# Phospho-KNL-1 recognition by a TPR domain targets the BUB-1–BUB-3 complex to *C. elegans* kinetochores

**DOI:** 10.1101/2024.02.09.579536

**Authors:** Jack Houston, Clémence Vissotsky, Amar Deep, Hiro Hakozaki, Enice Crews, Karen Oegema, Kevin D. Corbett, Pablo Lara-Gonzalez, Taekyung Kim, Arshad Desai

## Abstract

During mitosis, the Bub1-Bub3 complex concentrates at kinetochores, the microtubule-coupling interfaces on chromosomes, where it contributes to spindle checkpoint activation, kinetochore-spindle microtubule interactions, and protection of centromeric cohesion. Bub1 has a conserved N-terminal tetratricopeptide (TPR) domain followed by a binding motif for its conserved interactor Bub3. The current model for Bub1-Bub3 localization to kinetochores is that Bub3, along with its bound motif from Bub1, recognizes phosphorylated “MELT” motifs in the kinetochore scaffold protein Knl1. Motivated by the greater phenotypic severity of BUB-1 versus BUB-3 loss in *C. elegans*, we show that the BUB-1 TPR domain directly recognizes a distinct class of phosphorylated motifs in KNL-1 and that this interaction is essential for BUB-1–BUB-3 localization and function. BUB-3 recognition of phospho-MELT motifs additively contributes to drive super-stoichiometric accumulation of BUB-1–BUB-3 on its KNL-1 scaffold during mitotic entry. Bub1’s TPR domain interacts with Knl1 in other species, suggesting that collaboration of TPR-dependent and Bub3-dependent interfaces in Bub1-Bub3 localization and functions may be conserved.

## INTRODUCTION

During mitosis, kinetochores attach chromosomes to the mitotic spindle to enable accurate segregation of the replicated genome. To accomplish this, kinetochores establish dynamic microtubule attachments and continuously communicate with the core cell cycle engine. Originally discovered in yeast (Hoyt et al., 1991; Roberts et al., 1994), Bub1 is a conserved kinetochore component that is important for chromosome segregation across eukaryotes (Kim and Gartner, 2021). Bub1 has an N-terminal TPR domain, several short linear interaction motifs that interface with different chromosome segregation and regulatory factors, and a C-terminal kinase domain. Bub1 forms a complex with Bub3, a WD40 domain protein that binds to a short linear motif adjacent to the Bub1 TPR domain. Bub3 is implicated in the stability and kinetochore recruitment of Bub1 (Hoyt et al., 1991; Larsen et al., 2007; Roberts et al., 1994; Taylor et al., 1998). Bub3 is also present, without Bub1, in the mitotic checkpoint complex that unattached kinetochores produce to delay mitotic progression (Lara-Gonzalez et al., 2021b; McAinsh and Kops, 2023; Musacchio, 2015). Bub1 plays three important roles at kinetochores. First, Bub1 plays a central role in coordinating chromosome segregation with cell cycle progression. At unattached kinetochores, Bub1 recruits spindle checkpoint components and activates checkpoint signaling to delay mitotic progression (Di Fiore et al., 2015; Lara-Gonzalez et al., 2021a; London and Biggins, 2014; Moyle et al., 2014; Sharp-Baker and Chen, 2001). In specific contexts, such as the rapidly dividing *C. elegans* embryo, Bub1 also promotes mitotic progression by activating the Anaphase Promoting Complex/Cyclosome, the E3 ubiquitin ligase responsible for sister chromatid separation and mitotic exit (Kim et al., 2017; Kim et al., 2015). Second, Bub1 promotes centromeric cohesion by phosphorylating histone H2A and recruiting the Shugoshin/PP2A complex, which locally protects cohesin from removal prior to anaphase (Kawashima et al., 2010). Third, Bub1 plays an important role in chromosome alignment and segregation by contributing to the recruitment of components such as the chromosomal passenger complex, the dynein-recruiting RZZ complex, BubR1-PP2A (in vertebrates), and CENP-F (Ciossani et al., 2018; Edwards et al., 2018; Essex et al., 2009; Johnson et al., 2004; Klebig et al., 2009; Kruse et al., 2013; Overlack et al., 2015; Suijkerbuijk et al., 2012; Zhang et al., 2015). All of these functions rely on the targeting of Bub1 to kinetochores, highlighting the importance of understanding the dynamics and regulation of Bub1 recruitment to kinetochores.

The conserved N-terminal TPR domain of human Bub1 binds to a hydrophobic motif (referred to as KI1) in the kinetochore scaffold protein Knl1 (Bolanos-Garcia et al., 2009; Kiyomitsu et al., 2011; Krenn et al., 2012); the Bub1-related BubR1 protein similarly employs its TPR domain to interface with a second KI2 motif in Knl1 (Bolanos-Garcia et al., 2011; Kiyomitsu et al., 2011). Although the TPR-Bub1 interface has the potential to contribute to the kinetochore localization of Bub1, it remains unclear if it does so. When tested in nocodazole-treated HeLa cells, the Bub1-KI1 interface was found to be largely dispensable for Bub1 kinetochore localization (Krenn et al., 2012; Taylor et al., 1998). Evidence for KI-like motifs outside of vertebrate Knl1s is lacking and direct Bub1-Knl1 interactions, as observed for human Bub1 and Knl1, have not been reported. Consistent with pioneering work establishing the importance of Bub3 in Bub1 localization in human cells (Taylor et al., 1998), the molecular interface generally accepted as central to Bub1 kinetochore localization involves its Bub3 binding motif (also referred to as the GLEBS motif (Wang et al., 2001)), which is adjacent to the N-terminal TPR. Structural and biochemical analysis established that one side of the Bub3 WD40 barrel, along with the Bub3 binding motif of Bub1 that binds to the top of the WD40 barrel, recognize phosphorylated MELT repeats in Knl1 family proteins (Primorac et al., 2013). These repeats are phosphorylated by Mps1 and/or Plk1, depending on the species (Espeut et al., 2015; London et al., 2012; Shepperd et al., 2012; von Schubert et al., 2015; Yamagishi et al., 2012) and their phosphorylation is opposed by localized phosphatase activities, such as the PP1 that docks onto the N-terminus of Knl1 (London et al., 2012; Nijenhuis et al., 2014). Thus, the current model for Bub1 kinetochore localization posits that dynamic changes in phosphorylation/dephosphorylation of MELT motifs in Knl1 are read by a composite surface created by the Bub3 binding motif of Bub1 bound to Bub3; this composite surface is likely extended in specific organisms, as highlighted by the importance of a “SHT” motif following the MELT repeats in recognition by human Bub3 (Vleugel et al., 2015; Vleugel et al., 2013). While widely accepted, the Bub3-centric model for Bub1 localization leaves open the question as to why the Bub1 TPR domain is conserved across eukaryotes. Studies in human cells and fission yeast suggest that the Bub1 TPR contributes to the robustness of spindle checkpoint signaling (Klebig et al., 2009; Krenn et al., 2014; Leontiou et al., 2019). However, analysis of the Bub1 TPR has been relatively limited, leaving open other possible functions.

Here, we investigate the mechanisms that recruit BUB-1 to kinetochores in the *C. elegans* embryo. Our effort was inspired by the observation that BUB-1 depletion leads to a more severe defect in chromosome segregation than BUB-3 depletion or genetic deletion (Kim et al., 2015). We show that the TPR domain is essential for BUB-1 recruitment to kinetochores. The TPR domain recruits BUB-1 to kinetochores by recognizing a set of phosphorylated motifs in KNL-1 that are distinct from the MELT motifs. The ability of BUB-3 to recognize phospho-MELT motifs is required for the super-stoichiometric accumulation of the BUB-1–BUB-3 complex at kinetochores during mitotic entry but is dispensable for BUB-1’s essential functions. In addition to defining a TPR-centered mechanism for BUB-1 kinetochore localization, these results highlight the potential for TPR domains to function as phospho-readers, which may be of significance in contexts beyond chromosome segregation.

## RESULTS

### BUB-1 is required for chromosome segregation during mitosis independent of its prior roles in meiosis

The current model suggests that the Bub1-Bub3 complex is recruited to kinetochores via Bub3-mediated recognition of phosphorylated MELT motifs in the kinetochore scaffold protein Knl1 (**Fig. 1A**). This model predicts that loss of Bub3 should prevent Bub1 from being recruited to kinetochores and lead to chromosome segregation defects of similar severity to those following loss of Bub1. However, in *C. elegans,* the phenotypes associated with BUB-3 loss are significantly less severe than those resulting from BUB-1 loss (**Fig. 1A,B; Fig. S1A,B** (Essex et al., 2009; Kim et al., 2015)). RNAi-mediated depletion of BUB-1 leads to penetrant embryonic lethality, whereas BUB-3 depletion does not (**Fig. 1A**), a result consistent with prior analysis of null mutations in *bub-1* and *bub-3* (Kim et al., 2015). BUB-1 depletion also leads to significant chromosome missegregation in one-cell embryos whereas knockdown or mutation of BUB-3 does not (**Fig. 1B**; (Kim et al., 2015). Prior work has shown that the absence of BUB-3 leads to a significant (∼80%) reduction in BUB-1 protein levels (Kim et al., 2015). Collectively, these results suggest that the ∼20% of BUB-1 that remains in the absence of BUB-3 is sufficient to support chromosome segregation and viability, and raises the possibility that BUB-1 can localize to kinetochores and function independently of BUB-3 (Kim et al., 2015; Macaisne et al., 2023).

**Figure 1.**
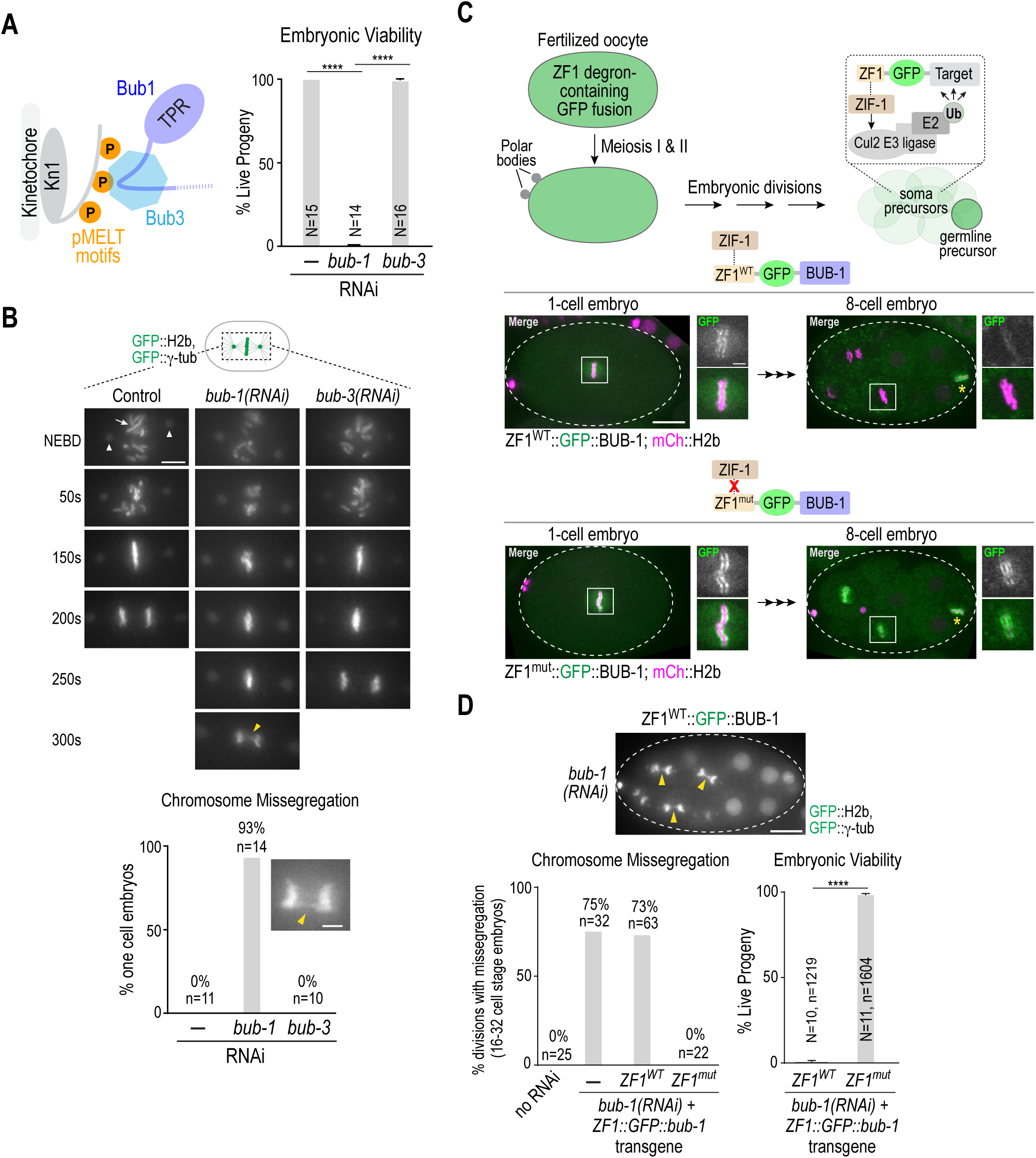
The more severe chromosome segregation defect of BUB-1 depletion relative to BUB-3 depletion is independent of BUB-1 function in oocyte meiosis. **(A)** (*left*) Current model for Bub1-Bub3 complex localization to kinetochores. (*right*) Embryo viability analysis for the indicated conditions. *N* is number of worms whose progeny were scored. Error bars are the 95% confidence interval. **(B)** (*top*) Stills from timelapse movies, aligned with NEBD, for the indicated conditions. Embryos imaged expressed GFP::H2b and GFP::γ-tubulin to visualize chromosomes (*arrow*) and spindle poles (*arrowheads*), respectively. Yellow arrowhead in the 300s *bub-1(RNAi)* panel highlights missegregation. Scale bar, 5 µm. (*bottom*) Quantification of chromosome missegregation in anaphase for the indicated conditions. Inset shows an example missegregation event. N is number of embryos imaged. Scale bar, 2 µm. **(C)** (*top*) Schematic describing the effect of ZF1 degron fusion to a target protein. The fusion is present during oocyte meiosis and in one-cell embryos but gets degraded specifically in soma precursor cells in a ZIF-1-dependent manner. (*bottom*) Images of ZF1^WT^ and ZF^mut^ GFP::BUB-1 fusions at 1-cell and 8-cell stages; the imaged embryos also expressed mCh::H2b to mark chromosomes. Dashed white line shows the embryo outline. White boxes indicate regions magnified on the right. Yellow asterisk marks the germline precursor cell. Scale bar, 10 µm (*whole embryo images*), and 2 µm (*magnified regions*). **(D)** (*top*) Image of GFP::H2b and GFP::γ-tubulin in a ZF1^WT^::GFP::BUB-1 embryo following depletion of endogenous BUB-1. Arrowheads point to missegregating chromosomes in soma precursor cells. Graphs below plot chromosome missegregation (monitored at the 16-32 cell stage, *left*) and embryo viability (*right*). Error bars are the 95% confidence interval. In the viability analysis *N* is the number of worms and *n* the number of progeny embryos scored. All *p* values are from unpaired t tests; **** = p<0.0001.

In *C. elegans*, BUB-1 is important for acentrosomal spindle assembly and chromosome segregation during oocyte meiosis, which occurs just prior to the first embryonic mitosis (Dumont et al., 2010; Macaisne et al., 2023), whereas depletion or genetic deletion of BUB-3 does not result in meiotic segregation defects (Macaisne et al., 2023). Thus, one possibility is that the mitotic segregation defects following BUB-1 depletion are a consequence of the prior defect in meiotic chromosome segregation, and BUB-3-depleted embryos do not exhibit similar mitotic defects because BUB-3 is not required for meiotic segregation. To address this possibility, we engineered RNAi-resistant transgenes encoding GFP::BUB-1 fused to the somatic cell-specific ZF1 degron recognized by the Cullin2 ubiquitin ligase adapter ZIF-1 (**Fig. 1C**; (Armenti et al., 2014; Reese et al., 2000)). The ZF1 degron is derived from the germline determinant PIE-1, which is maternally loaded but then selectively degraded by the 8-16 cell stage in somatic tissue precursor cells in a ZIF-1-dependent manner (**Fig. 1C**; (Reese et al., 2000)). A ZF1 degron containing point mutations that prevent ZIF-1 binding and target degradation (ZF1^mut^) served as a control. GFP::BUB-1 fusions with ZF1^WT^ and ZF1^mut^ were both expressed during oocyte meiosis and in 1- and 2-cell embryos, after which the ZF1^WT^ fusion was degraded in developing somatic cells whereas the ZF1^mut^ fusion was not (**Fig. 1C**). Following depletion of endogenous BUB-1, embryos expressing ZF1^WT^-GFP::BUB-1 exhibited chromosome segregation defects during somatic precursor cell divisions after the fusion was degraded and penetrant lethality, whereas embryos expressing the non-degradable ZF1^mut^-GFP::BUB-1 fusion did not (**Fig. 1D**). We conclude that the more severe mitotic phenotypes in BUB-1 compared to BUB-3 depleted embryos are not explained by the differential roles of the two proteins in meiosis. Instead, it is likely that mitotic phenotypes are less severe in BUB-3 depleted embryos because a BUB-1 pool is present and retains the ability to localize to kinetochores and function independently of BUB-3.

### The BUB-1 TPR domain is necessary and sufficient for kinetochore localization and is required for chromosome segregation

The results above suggest that BUB-1 can localize to kinetochores independently of BUB-3. To identify functional elements in BUB-1, independent from its well-defined BUB-3 binding motif, that are sufficient for kinetochore recruitment we employed an RNAi-resistant transgene system that enables replacing endogenous BUB-1 with engineered variants (Moyle et al., 2014).

Work in human cells has suggested that the Bub1 TPR contributes to its kinetochore localization (Kiyomitsu et al., 2011; Kiyomitsu et al., 2007; Klebig et al., 2009) and subsequent work revealed the structure of the Bub1 TPR bound to a short peptide motif, named KI1, in Knl1 (Krenn et al., 2012). However, cell biological analysis conducted in parallel with the structural work, in agreement with the first study on human Bub3 (Taylor et al., 1998), argued against the Bub1 TPR making a substantial contribution to Bub1 kinetochore localization (Krenn et al., 2012). Despite differing conclusions, these prior studies suggested that a TPR-mediated interaction with KNL-1 could be a potential mechanism for BUB-3-independent localization of BUB-1 in *C. elegans*.

To address the possibility that the TPR is important for BUB-1 kinetochore localization, we generated strains harboring single copy RNAi-resistant transgenes expressing GFP fusions with WT BUB-1, BUB-1 lacking the TPR domain (ΔTPR), or the TPR domain on its own (TPR only) and confirmed expression by immunoblotting (**Fig. 2A,B**). Following endogenous BUB-1 depletion, the GFP fusions with WT BUB-1 or the BUB-1 TPR domain alone localized to kinetochores, whereas the fusion with BUB-1 lacking the TPR domain failed to be recruited (**Fig 2C**). Quantification revealed that the TPR-only BUB-1 fragment was recruited to kinetochores at ∼40% of the level of wild-type BUB-1 (**Fig. 2C**). Thus, the TPR domain is both necessary and sufficient to recruit BUB-1 to kinetochores, albeit at a reduced level compared to WT BUB-1. We note that ΔTPR BUB-1 failed to localize to kinetochores even in the presence of endogenous BUB-1 **(Fig. S1C)**, which is consistent with kinetochore recruitment requiring a direct interaction between the TPR domains of individual BUB-1 molecules and the kinetochore scaffold.

**Figure 2:**
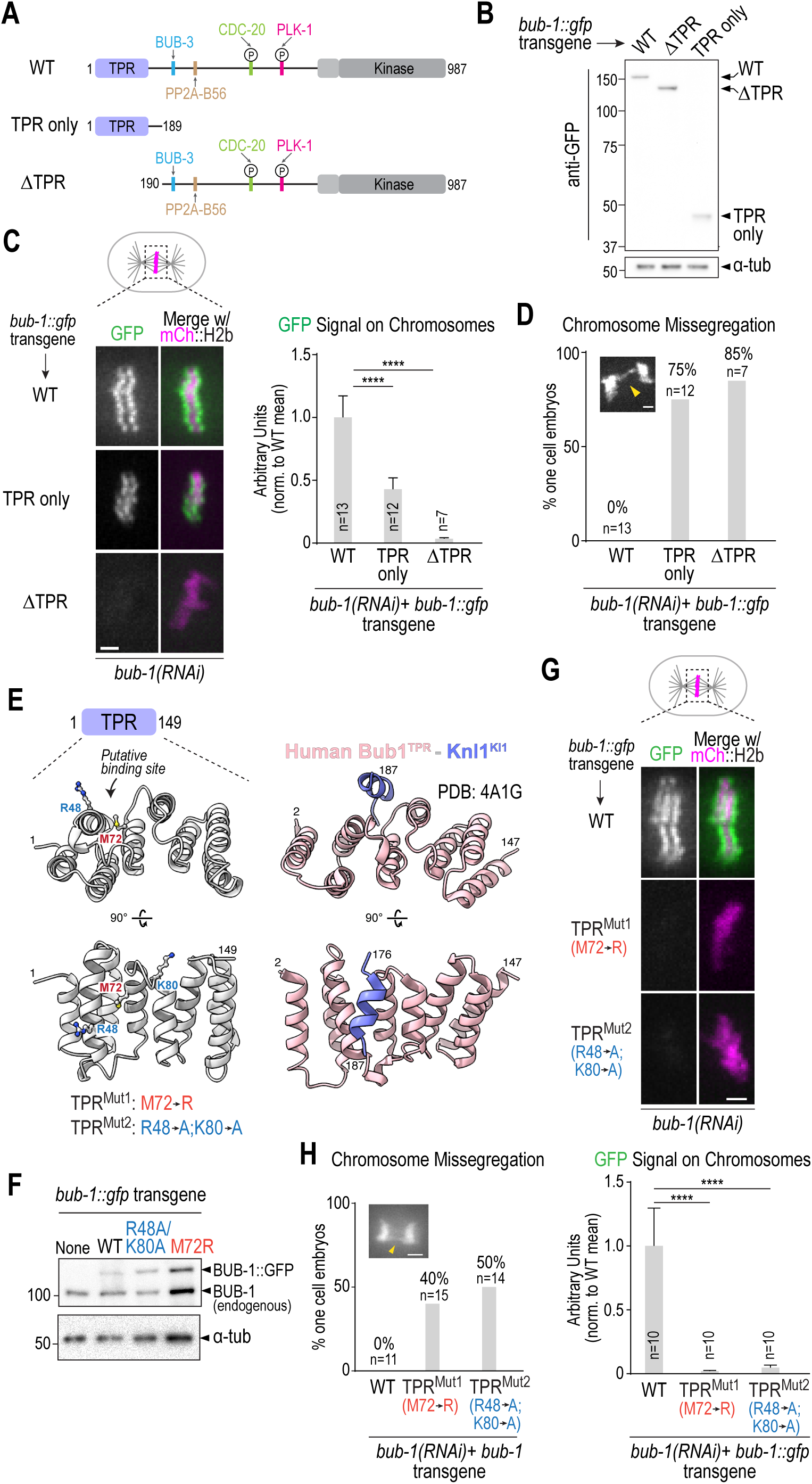
The BUB-1 TPR is necessary and sufficient for kinetochore localization. **(A)** Schematic of wild-type BUB-1, highlighting key domains and interfaces, along with designed TPR only and ΔTPR variants. **(B)** Anti-GFP immunoblots of strains with the indicated integrated *bub-1::gfp* transgenes. Molecular weight markers are in kilodaltons. α-tubulin serves as a loading control. **(C)** (*left*) Images of aligned chromosomes in one-cell embryos from strains with the indicated *bub-1::gfp* transgenes; the strains also expressed mCh::H2b to mark chromosomes and endogenous BUB-1 was depleted. Scale bar, 2 µm. (*right*) Quantification of GFP signal on chromosomes for the indicated conditions. Error bars are the 95% confidence interval. *n* is the number of embryos imaged. **(D)** Quantification of chromosome missegregation in one-cell embryo anaphase for the indicated conditions. *n* is the number of embryos imaged. **(E)** Comparison of a structural model of the BUB-1 TPR and the human Bub1 TPR – Knl1 KI1 motif crystal structure (PDB:4A1G) (Krenn et al., 2012). Residues targeted for mutation in BUB-1 are highlighted below the *en face* view of the convex surface of the TPR. **(F)** Anti-BUB-1 immunoblots of strains with the indicated integrated *bub-1::gfp* transgenes. GFP-fused and endogenous BUB-1 are marked. α-tubulin serves as a loading control. **(G)** (*top*) Images of aligned chromosomes in one-cell embryos from strains with the indicated *bub-1::gfp* transgenes; the strains also expressed mCh::H2b to mark chromosomes and endogenous BUB-1 was depleted. Scale bar, 2 µm. (*bottom*) Quantification of GFP signal on chromosomes for the indicated conditions. *n* is the number of embryos imaged. Error bars are the 95% confidence interval. **(H)** Quantification of chromosome missegregation in one-cell embryos for the indicated conditions. *n* is the number of embryos imaged. Example missegregation image is the same as in Fig. 1B. All *p* values were calculated by unpaired t tests; **** = p<0.0001.

Following endogenous BUB-1 depletion, ΔTPR BUB-1, which fails to localize to kinetochores, exhibited chromosome segregation defects comparable to those observed following BUB-1 depletion (**Fig. 2D**). The TPR-only BUB-1 fragment also failed to support chromosome segregation (**Fig. 2D**), which is expected since it lacks all other interaction surfaces and functional domains of BUB-1. Collectively, these data show that the TPR domain is critical for the kinetochore localization and functions of BUB-1 in the *C. elegans* embryo.

### Structure-guided mutations in the BUB-1 TPR domain disrupt BUB-1 localization and function

While *C. elegans* KNL-1 lacks a clear KI-like motif found in human Knl1, we suspected that it might interact with the *C. elegans* BUB-1 TPR in a manner similar to that described for the human proteins. Comparing the crystal structure of human Bub1 TPR bound to the Knl1 KI1 motif (PDB:4A1G: (Krenn et al., 2012)) to a structural model of the *C. elegans* BUB-1 TPR domain led us to design two mutants (M72R and R48A/K80A) that we predicted would disrupt interaction of the *C. elegans* BUB-1 TPR with a KI-like motif (**Fig. 2E**). We generated strains with single-copy integrated transgenes encoding these mutant forms of BUB-1, confirmed their expression (**Fig. 2F**), and analyzed their localization in the absence of endogenous BUB-1 (**Fig. 2G**). Both mutations nearly completely eliminated BUB-1 kinetochore localization, suggesting that an interaction with the convex surface of the BUB-1 TPR, analogous to that observed with human Bub1 TPR-Knl1 KI1 (Krenn et al., 2012), is critical for BUB-1 localization in the *C. elegans* embryo. As predicted by the localization defect, both mutants exhibited chromosome missegregation and embryonic lethality in the absence of endogenous BUB-1 (**Fig. 2H; Fig. S1D**). Both mutants also failed to localize in the presence of endogenous BUB-1 (**Fig. S1E**), as was also observed with ΔTPR BUB-1.

Collectively, the above data indicate that the BUB-1 TPR domain is both necessary and sufficient for kinetochore localization and that it employs an interface on its convex surface, analogous to the one defined for human Bub1–Knl1 KI1, to localize and function at the kinetochore.

### Recruitment of BUB-1 to kinetochores requires putative TPR-interacting motifs in KNL-1

To identify potential KI-like motifs in KNL-1 we turned to AlphaFold predictions (Evans et al., 2022). AlphaFold predicted with medium-to-high confidence that the BUB-1 TPR domain interfaces with repetitive motifs in the KNL-1 N-terminus that have the consensus [D/N]-[D/E]-T-[<]-[x]-[<]-F, where the T and F are invariant (**Fig. 3A,B; Fig. S2**). This same motif, which is distinct from KI motifs, was identified in prior work by phylogenetic sequence analysis as being present in nematode and insect Knl1s (Tromer et al., 2015), and we refer to it as the TF motif (**Fig. 3A, Fig. S3A**). Interestingly, the TF motif shares similarity to the TΩ motif that precedes the MELT motifs in human Knl1 that are active for Bub1-Bub3 recruitment (Vleugel et al., 2015). There are six TF motifs in the *C. elegans* KNL-1 N-terminus. In our AlphaFold models, the phenylalanine of five of the six TF motifs docks into the same hydrophobic pocket formed on the convex surface of the TPR domain by alpha helices 3-5 (**Fig. 3B; Fig. S2**). Within the pocket, the TF motifs are in contact with the Met72 residue of BUB-1, which may explain why the Met72Arg mutation has such a strong effect on kinetochore localization.

**Figure 3.**
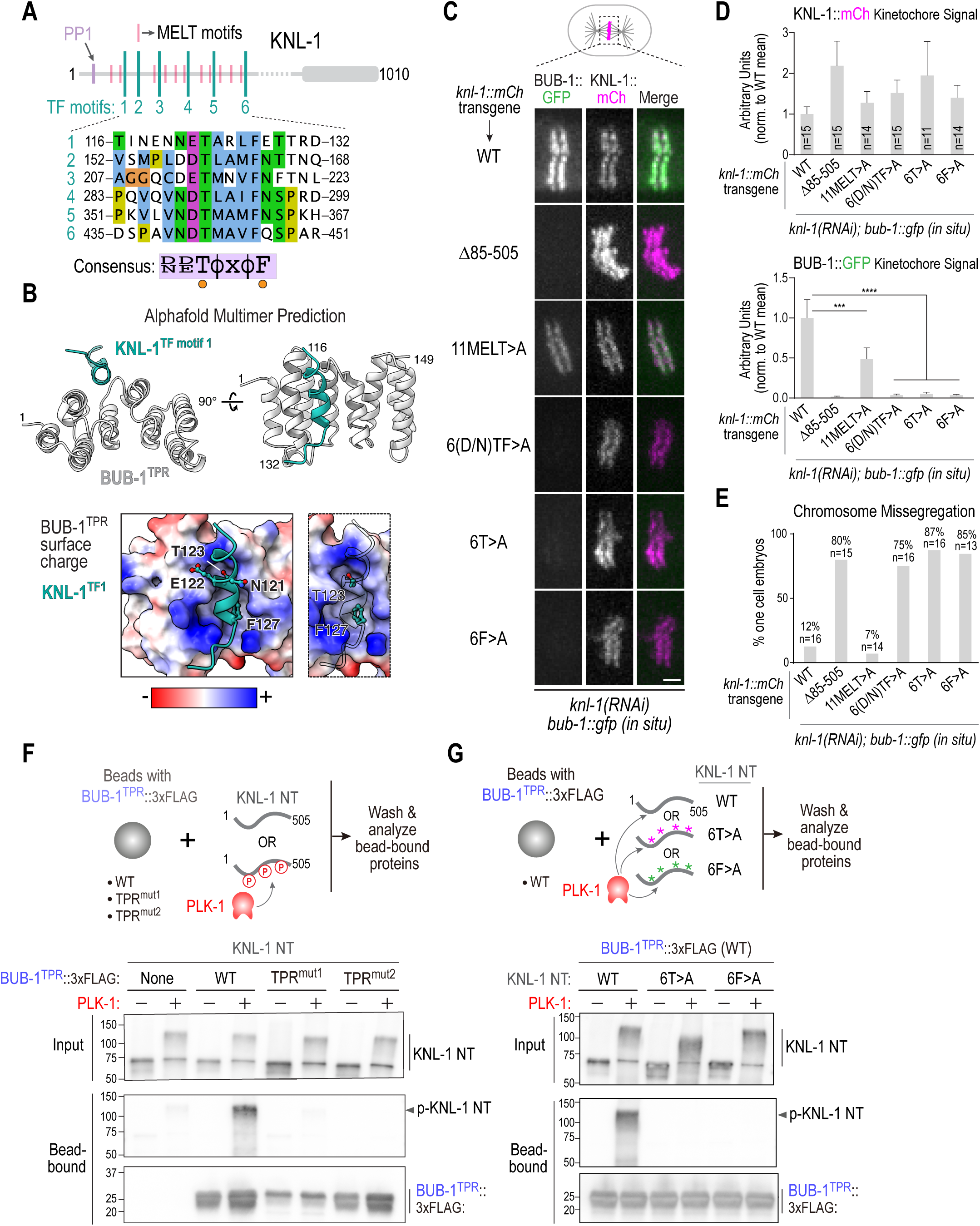
Definition of the BUB-1 TPR – KNL-1 interface that targets BUB-1 to kinetochores. **(A)** Schematic of KNL-1 highlighting 11 MELT motifs, 6 TF motifs and protein phosphatase 1 (PP1) docking site. Sequences of the 6 TF motifs are shown below along with a consensus. **(B)** (*top*) AlphaFold multimer prediction of the BUB-1 TPR interface with TF motif 1. Side and *en face* views are shown. Predictions for all 6 TF motifs are shown in *Fig. S2*. (*bottom*) TF motif 1 shown docked onto a surface charge model of the BUB-1 TPR. Panel on the right highlights the T and F residues; the insertion of the F into a hydrophobic pocket is evident in this view. **(C)** (*left*) Images of aligned chromosomes in one-cell embryos from strains with the indicated *knl-1::mCh* transgenes and *in situ* GFP-tagged BUB-1; endogenous KNL-1 was depleted in all conditions. Scale bar, 2 µm. **(D)** Quantification of KNL-1::mCh and BUB-1::GFP kinetochore signals for the indicated conditions. *n* is the number of embryos imaged. Error bars are the 95% confidence interval. **(E)** Quantification of chromosome missegregation in one-cell embryos for the indicated conditions. *n* is the number of embryos imaged. **(F)** (*top*) Schematic of biochemical assay used to assess BUB-1 TPR interaction with the KNL-1 N-terminus, either with or without PLK-1 phosphorylation. The BUB-1 TPR variants were expressed in suspension human cells and concentrated on beads prior to mixing with unphosphorylated or PLK-1 phosphorylated bacterially expressed and purified KNL-1 N-terminus (NT). (*bottom*) Immunoblots of the KNL-1 input and bead-bound KNL-1 and BUB-1^TPR^ variants. **(G)** Similar biochemical analysis as in *(F)*, except that BUB-1^TPR^ was WT in all conditions and three versions of recombinant KNL-1 NT (WT, 6T>A, 6F>A) were tested with and without PLK-1 phosphorylation. Data shown in *(F)* and *(G)* is representative of two independent experiments. All *p* values were calculated by unpaired t tests; *** = p<0.001,**** = p<0.0001.

To determine if the TF motifs in KNL-1 predicted by AlphaFold to interface with the BUB-1 TPR are important for BUB-1 kinetochore localization, we employed a transgene-based KNL-1 replacement system (Espeut et al., 2012) to engineer three mutants: “6(D/N)TF>A”, in which all of the (D/N), T, and F residues in the motifs were mutated to alanines; “6T>A”, in which the threonines in the 6 motifs were mutated to alanines; and “6F>A”, in which the phenylalanines of the 6 motifs were mutated to alanines. In addition, we generated a transgene that mutated the threonines in 11 “MELT” motifs of KNL-1 to alanines, which we termed the “11MELT>A” mutant (**Fig. 3C**). We crossed transgenes expressing these KNL-1 variants, along with transgenes expressing WT KNL-1 and a deletion of the majority of the N-terminal half of KNL-1 that removes all MELT and TF motifs (Δ85-505; (Moyle et al., 2014)) as positive and negative controls, into a strain in which BUB-1 was tagged at its endogenous locus with GFP (**Fig. 3C**). Following depletion of endogenous KNL-1, we monitored BUB-1 and KNL-1 kinetochore localization. None of the KNL-1 variants significantly reduced the ability of KNL-1 to localize to kinetochores (**Fig. 3C,D**). As expected, transgene-encoded WT KNL-1 supported robust BUB-1 localization, whereas Δ85-505 KNL-1, which removes all of the MELT and TF motifs, exhibited no detectable BUB-1 localization (**Fig. 3C,D**). All three TF motif mutants significantly compromised BUB-1 localization (**Fig. 3C,D**); by contrast, the 11MELT>A mutant only modestly affected BUB-1 localization (**Fig. 3C,D**). In the strain with endogenously tagged BUB-1, the KNL-1 TF motif mutants exhibited chromosome missegregation and embryonic lethality while the 11MELT>A mutant had a much milder effect (**Fig. 3E**; **Fig. S3B**). In a strain background in which the endogenous *bub-1* locus was not tagged, the TF mutants exhibited missegregation and lethality but the phenotypic penetrance was less severe (**Fig. S3C,D**), which is indicative of a negative synthetic genetic interaction between TF motif mutants of KNL-1 and *in situ* GFP-tagged BUB-1.

Collectively, the structural modeling and *in vivo* analysis support a model in which the TPR domain of BUB-1 engages with TF motifs in the KNL-1 N-terminus to target BUB-1 to kinetochores.

### The BUB-1 TPR binds to the KNL-1 N-terminus in a PLK-1 phosphorylation-dependent manner

While the AlphaFold modeling revealed a potential direct interface between TF motifs of KNL-1 and the BUB-1 TPR, biochemical and 2-hybrid efforts over multiple years had failed to detect an interaction between the KNL-1 N-terminus and BUB-1, which could be due to a lack of required phosphoregulation. We noticed that the threonines in the KNL-1 TF motifs perfectly match a Polo-like Kinase 1 (PLK-1) consensus phosphorylation site (Santamaria et al., 2011). We therefore speculated that PLK-1 phosphorylates TF motif threonines to enable interaction with the BUB-1 TPR. To test this hypothesis, we used a biochemical assay to analyze interaction between the BUB-1 TPR and the KNL-1 N-terminus, with and without PLK-1 phosphorylation (**Fig. 3F**). WT or mutant variants of the BUB-1 TPR that impaired localization and function *in vivo* were expressed in human suspension cell culture and concentrated on beads. Bacterially-expressed KNL-1 N-terminus, with or without PLK-1 phosphorylation, was incubated with the BUB-1 TPR variant-coated beads and the beads were isolated and analyzed by immunoblotting (**Fig. 3F**). The purified KNL-1 N-terminus bound robustly to the WT BUB-1 TPR beads when it was phosphorylated by PLK1, whereas no significant binding was observed in the absence of phosphorylation (**Fig. 3F**). The BUB-1 TPR mutants that disrupted kinetochore localization *in vivo* also failed to bind to the phosphorylated KNL-1 N-terminus *in vitro* (**Fig. 3F**). On the KNL-1 side, introducing the 6T>A or the 6F>A mutations into the KNL-1 N-terminus also prevented it from binding to the BUB-1 TPR (**Fig. 3G**). Both mass spectrometry (**Fig. S3E**) and reduced retardation of electrophoretic mobility (**Fig. 3G**), provided evidence for phosphorylation of TF motifs by PLK-1 (**Fig. 3G**).

Taken together, the structural modeling, *in vivo* characterization of engineered mutants, and biochemical analysis support the model that the BUB-1 TPR recruits BUB-1 to kinetochores by directly engaging PLK-1 phosphorylated TF motifs in the KNL-1 N-terminus.

### BUB-3-dependent MELT recognition drives the super-stoichiometric recruitment of BUB-1–BUB-3 complexes relative to their kinetochore scaffold during mitotic entry

Next, we wanted to address the role of BUB-3, which has the ability to directly bind to phosphorylated KNL-1 MELT (pMELT) repeats (London et al., 2012; Primorac et al., 2013; Shepperd et al., 2012; Yamagishi et al., 2012). Since BUB-1 TPR mutants prevented recruitment of BUB-1 to kinetochores, we first tested whether they also prevented kinetochore recruitment of BUB-3. This effort revealed that disruption of the BUB-1 TPR-TF motif interface led to the near-complete elimination of kinetochore-localized BUB-3 (**Fig. 4A**). Thus, free BUB-3 is not recruited to kinetochores, and BUB-3-dependent pMELT recognition cannot recruit the BUB-1–BUB-3 complex to kinetochores in the absence of phospho-TF motif recognition by the BUB-1 TPR domain.

**Figure 4.**
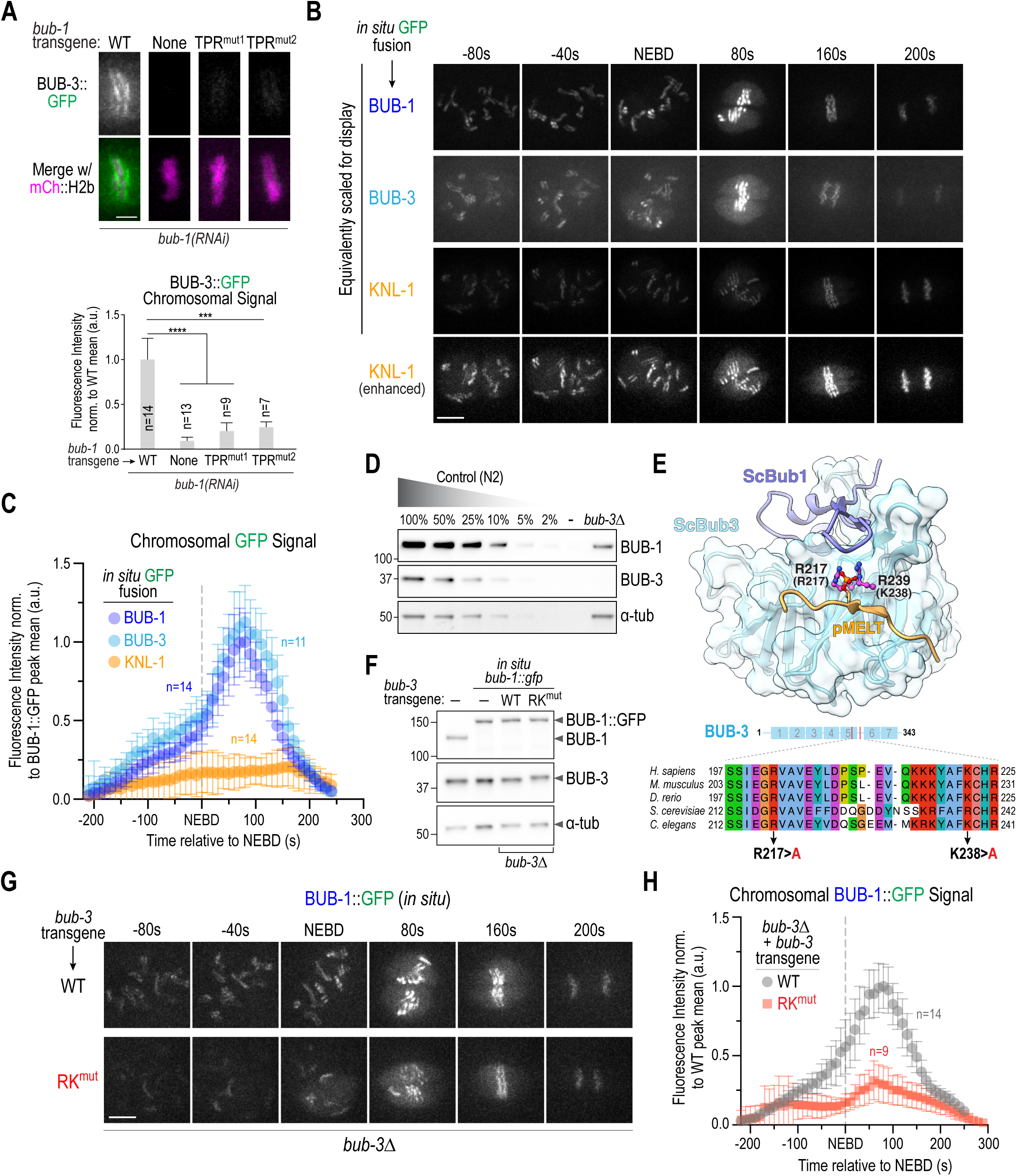
BUB-3 recognition of pMELT motifs drives the rapid increase in BUB-1–BUB-3 localization at kinetochores during mitotic entry. **(A)** (*top*) Localization of BUB-3::GFP in the indicated conditions. Scale bar, 2 µm. (*bottom*) Quantification of BUB-3::GFP chromosomal signal. Error bars are the 95% confidence interval. **(B)** Images from timelapse sequences of the indicated *in situ* GFP fusions. Sequences were time-aligned using NEBD as a reference. All three strains also expressed mCh::H2b which was simultaneously imaged but is not shown. All imaging was conducted under identical conditions; the top 3 image rows are equivalently scaled for display to highlight the super-stoichiometric recruitment of BUB-1–BUB-3 complexes on KNL-1 scaffolds. The bottom row shows enhanced scaling of the KNL-1::GFP signal to display its robust and constant kinetochore localization. Scale bar, 5 µm. **(C)** Chromosome segmentation-based quantification of GFP signal over time for the indicated *in situ* GFP fusions. Chromosome segmentation was performed using the mCh::H2b signal (see *Fig. S4A* for details). Error bars are the 95% confidence interval. *n* is the number of embryos imaged. **(D)** Immunoblots of a dilution curve of control (N2) worm extract and a *bub-3Δ* worm extract using anti-BUB-1 and anti-BUB-3 antibodies. α-tubulin serves as a loading control. **(E)** (*top*) Structure of *S. cerevisiae* Bub1-Bub3 complex bound to a pMELT peptide from ScKnl1/Spc105 (PDB:4BL0;(Primorac et al., 2013)). Sidechains of the phosphorylated Thr in the MELT peptide and the two basic residues critical for binding the phosphopeptides (R217 and K239) are shown. Residue numbers in brackets below the R217 and K239 labels refer to the corresponding *C. elegans* BUB-3 residues. (*bottom*) Sequence alignment of the region of Bub3 that engages the pMELT peptide. The two basic residues mutated in BUB-3 to disrupt pMELT recognition are highlighted. **(F)** Immunoblots of the indicated conditions using anti-BUB-1 and anti-BUB-3 antibodies. α-tubulin serves as a loading control. **(G)** Images from timelapse sequences of *in situ* GFP-tagged BUB-1 in the presence of WT or RK^mut^ BUB-3; the endogenous *bub-3* gene was deleted. The strains also expressed mCh::H2b, which was simultaneously imaged but is not shown. Scale bar, 5 µm. **(H)** Chromosome segmentation-based quantification of BUB-1::GFP signal over time for the indicated conditions. *n* is the number of embryos imaged. Error bars are the 95% confidence interval. All *p* values are calculated from unpaired t tests; *** = p<0.001,**** = p<0.0001.

While the BUB-1 TPR domain is sufficient to localize to kinetochores, it does not accumulate to the same extent as wildtype BUB-1 (**Fig. 2C**) suggesting that BUB-3-dependent pMELT recognition might synergize with the TPR-based interface to enhance kinetochore recruitment of the BUB-1–BUB-3 complex. To address the role of BUB-3-dependent pMELT recognition, we first developed a means to quantitatively compare the localization dynamics of BUB-1, BUB-3 and KNL-1 by imaging *in situ* GFP fusions for all three under identical conditions (**Fig. 4B**). This effort revealed a significant increase in BUB-1 and BUB-3 signal at kinetochores around the time of NEBD, which was not observed for KNL-1 and is indicative of super-stoichiometric association of BUB-1–BUB-3 complexes on the KNL-1 scaffold that recruits them to kinetochores. Using an image segmentation-based analysis approach (**Fig. S4A**), quantification of chromosomal GFP signals revealed peak accumulation of ∼five BUB-1 and BUB-3 molecules per KNL-1 molecule at kinetochores (**Fig. 4C**). Thus, one attractive hypothesis is that BUB-3 bound to BUB-1 contributes to super-stoichiometric accumulation of the complex at kinetochores that is coupled to mitotic entry.

To address the contribution of BUB-3 to BUB-1 kinetochore localization, we could not analyze BUB-3 deletion or depletion as they reduce BUB-1 protein levels by ∼80% (**Fig 4D**; (Kim et al., 2015)). Thus, we capitalized on prior structural work, along with the high sequence conservation of BUB-3, to engineer two point mutations (R217A and K238A: RK^mut^) that are expected to specifically disrupt pMELT recognition (**Fig. 4E**). To compare wildtype to RK^mut^ BUB-3 we built a transgene-based replacement system for BUB-3 (**Fig. S4B**). RK^mut^ BUB-3, unlike BUB-3 deletion or depletion, did not reduce BUB-1 protein levels (**Fig. 4F**). RK^mut^ BUB-3 significantly reduced BUB-1 localization and quantitative analysis showed that it largely prevented the super-stoichiometric accumulation of BUB-1–BUB-3 complex relative to KNL-1 observed when WT BUB-3 was present (**Fig. 4G,H**). These data suggest that, while a TPR-dependent mechanism is essential to localize the BUB-1–BUB-3 complex to kinetochores, BUB-3-dependent pMELT recognition is required to drive super-stoichiometric recruitment.

### BUB-1-bound PLK-1 contributes to the super-stoichiometric recruitment of BUB-1 to kinetochores during mitotic entry

In *C. elegans*, the kinase that targets both the TF motifs and MELT repeats in the N-terminus of KNL-1 is PLK-1 ((Espeut et al., 2015); **Fig. 3F,G**). There are at least 2 pools of PLK-1 docked at *C. elegans* kinetochores: one bound to BUB-1 and the second bound to CENP-C^HCP-4^ (Taylor et al., 2023). We had previously engineered a point mutation in the PLK-1 docking motif of BUB-1 (BUB-1 PD^mut^) and observed that it reduced but did not eliminate BUB-1 localization at kinetochores (Houston et al., 2023). Live imaging followed by segmentation-based quantitative analysis of transgene-encoded GFP fusions of wildtype and PD^mut^ BUB-1 revealed that the increase in BUB-1 localization at kinetochores was significantly suppressed when PLK-1 docking to BUB-1 was prevented (**Fig. 5A,B**). Because PD^mut^ BUB-1 exhibits embryonic lethality following depletion of endogenous BUB-1, this analysis required the use of transgene-encoded BUB-1::GFP fusions. As prior analysis employed *in situ* GFP-tagged BUB-1 and different imaging conditions (**Fig. 4B,C,G,H**), the localization profiles across these experiments are not quantitatively comparable. Nonetheless, these data support an important role for BUB-1-docked PLK-1 in driving the super-stoichiometric recruitment of the BUB-1–BUB-3 complex to kinetochores during mitotic entry.

**Figure 5.**
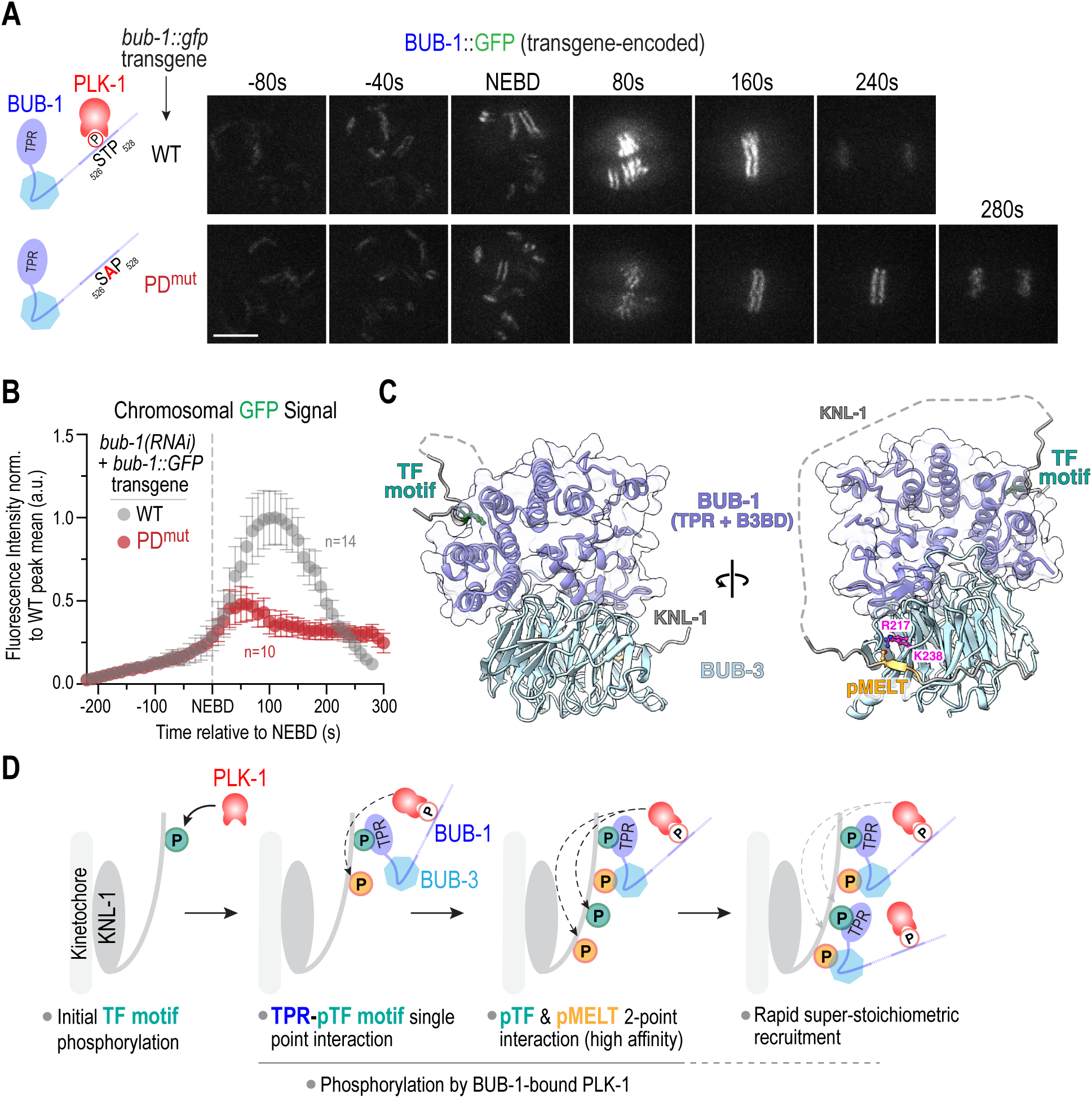
PLK-1 docked on BUB-1 is required for the increase in BUB-1–BUB-3 localization at kinetochores during mitotic entry. **(A)** (*left*) Schematics of WT BUB-1, which has a conserved Polo-like kinase 1 (PLK-1) docking site, and PD^mut^ BUB-1, in which T527 that, when phosphorylated, engages the Polo box domain of PLK-1 is mutated to an alanine. (*right*) Images from timelapse series of transgene-encoded WT and PD^mut^ BUB-1; mCh::H2b images were acquired at the same timepoints but are not shown. Scale bar, 5 µm. **(B)** Segmentation-based quantification of chromosomal localization over time for transgene-encoded WT and PD^mut^ BUB-1. *n* is the number of embryos imaged. Error bars are the 95% confidence interval. **(C)** AlphaFold multimer model of the BUB-1 N-terminus (TPR + BUB-3 Binding Domain), BUB-3, TF1 and MELT1 motifs from KNL-1. The TF motif interface is spatially distant from the pMELT-binding region, which enables formation of a higher-affinity two-point interface. **(D)** Model highlighting the steps leading to rapid, super-stoichiometric recruitment of BUB-1–BUB-3 complexes on a KNL-1 scaffold at kinetochores. See text for details.

## DISCUSSION

Here we provide a mechanistic resolution to the puzzling observation that loss of BUB-1 is far more severe than loss of BUB-3 in *C. elegans*, even though BUB-3-dependent recognition of pMELT repeats in the kinetochore scaffold protein KNL-1 was considered to be the primary mechanism that recruits the BUB-1–BUB-3 complex to kinetochores. Our efforts revealed that the convex surface of the BUB-1 TPR engages with specific phosphorylated TF motifs in the KNL-1 N-terminus in a manner reminiscent of the Bub1/BubR1 TPR – KI motif interaction in humans (Bolanos-Garcia et al., 2011; Krenn et al., 2012). This TPR-based interface is essential for BUB-1 recruitment and function at *C. elegans* kinetochores. We further show that while the TPR-dependent interface is essential to recruit the BUB-1–BUB-3 complex to kinetochores, BUB-3-dependent pMELT recognition is required for the super-stoichiometric recruitment of the BUB-1–BUB-3 complexes on KNL-1 scaffolds. Prior structures from other species and structural modeling of the *C. elegans* proteins supports the ability for both interfaces to co-exist and potentially generate a higher-affinity bound state (**Fig. 5C**, **5D**). Both interfaces are phosphoregulated, with PLK-1 being the kinase that targets the TF and MELT motifs. Analysis of a mutant form of BUB-1 that selectively removes its associated PLK-1 suggests that this specific pool of PLK-1 targets the MELT motifs that are engaged in a BUB-3-dependent manner. These results lead to a model in which a nucleating phosphorylation event of one or more TF motifs, by a pool of PLK-1 distinct from that bound to BUB-1, leads to engagement with the BUB-1 TPR (**Fig. 5D**). This in turn brings associated PLK-1 kinase activity that phosphorylates MELT motifs in the KNL-1 N-terminus, and possibly also additional TF motifs, to generate stable two-point interactions and rapidly drive super-stoichiometric association of BUB-1–BUB-3 complexes on KNL-1 scaffolds (**Fig. 5D**). The stoichiometry of BUB-1–BUB-3 complexes recruited relative to KNL-1 appears to match well the number of TF motifs, which supports the central importance of the TPR–TF motif interface. As both interfaces are phospho-dependent, they would be rapidly reversible by the action of localized (e.g. KNL-1-bound PP1) and global phosphatase activities. The pool of PLK-1 involved in the nucleating TF motif phosphorylation remains to be defined – it may either be nucleoplasmic active PLK-1 or the PLK-1 that associates with CENP-C^HCP-4^.

While disrupting the TPR–TF motif interface leads to near-complete loss of BUB-1 localization and penetrant chromosome missegregation in mitotically dividing embryonic cells, preventing the increase in BUB-1–BUB-3 localization, for example by selectively disrupting pMELT recognition, does not lead to embryonic lethality or obvious chromosome segregation defects (**Fig. S4C**)(Espeut et al., 2015). However, it does prolong mitotic duration (**Fig S4D**), likely by reducing kinetochore-dependent dephosphorylation and activation of CDC-20 (Houston et al., 2023; Kim et al., 2017), which our prior work has shown to impact the robustness of embryogenesis (Houston et al., 2023). The increase in BUB-1– BUB-3 localization at kinetochores is likely also critical for spindle checkpoint signaling, where the multiplicity of recruitment of Bub1-Bub3 complexes on Knl1 scaffolds is important (Chen et al., 2019; Vleugel et al., 2015; Zhang et al., 2014).

The conclusions from our analysis in *C. elegans* are distinct from the current view of how Bub1-Bub3 localization operates in human cells and budding yeast. While some studies in human cells have suggested a role for the TPR in Bub1 localization (Kiyomitsu et al., 2011; Klebig et al., 2009), other studies concluded that Bub1 localization was independent of the TPR and primarily dependent on the Bub3-binding domain (Krenn et al., 2012; Taylor et al., 1998). The latter view is currently widely accepted for how Bub1-Bub3 complexes localize to kinetochores in human cells. However, in both human cells and *C. elegans*, the Bub1 TPR domains employ their convex surface to directly engage with the N-termini of their respective Knl1s. This similarity across a wide evolutionary distance suggests an ancient and direct connection between the Bub1 TPR and Knl1. To date, precise mutations disrupting TPR-Knl1 interactions have only been analyzed with truncated forms of Knl1 in human cells, which indicated an importance for this interaction in the robustness of spindle checkpoint signaling (Krenn et al., 2014). We note that other key kinetochore-localized regulators, including Mps1 kinase that is absent in *C. elegans* and BubR1 pseudokinase/Mad3s, have a TPR domain in many species, and that the MELT repeats that are active for Bub1-Bub3 recruitment in human Knl1 have associated TΩ motifs that share similarity to the TF motifs we analyze here (Tromer et al., 2016; Tromer et al., 2015; Vleugel et al., 2015). Thus, TPR-based recognition of motifs in Knl1 that are distinct from the phospho-MELT repeats recognized by Bub3 family proteins may be widespread and utilized in different ways in different species/contexts.

The TPR domain was first defined over 30 years ago in a gene that, interestingly, was later found to encode a core subunit of the anaphase-promoting complex, a central player in chromosome segregation (Sikorski et al., 1990). TPR domains are widespread and found in close to 1000 human proteins that are involved in myriad cellular functions. Engagement of the concave surface of the helical TPR domain with a peptide ligand is a commonly observed interaction mode, but other interaction types are also observed (Perez-Riba and Itzhaki, 2019). Both prior work and the current study indicate that the Bub1 TPR is unusual in that it employs its convex surface to engage peptide ligands in Knl1 family proteins. In addition, the TPR in *C. elegans* BUB-1 binds to its peptide ligand in a phosphorylation-dependent manner, which is a mode of TPR–ligand interaction that, to our knowledge, has not been previously reported. Thus, in addition to addressing the mechanism of BUB-1–BUB-3 kinetochore localization across species, the findings here suggest the potential for TPR domains to act as phospho-readers, which may be relevant in contexts beyond chromosome segregation.

## ACKNOWLEDGEMENTS

The authors thank Eelco Tromer (University of Gröningen) and members of the Oegema and Desai labs for helpful discussions. We thank Andreas Ernst and Savannah Bogus (UC San Diego) for help culturing HEK293F Freestyle cells, and Jeffrey Woodruff (UT Southwestern) for constitutively active PLK-1. Mass spectrometry analysis of KNL-1 phospho-sites was conducted by Majid Ghassemian at the UCSD Biomolecular and Proteomics Mass Spectrometry Facility (NIH shared instrumentation grant numbers S10 OD021724). This work was supported by grants from the NIH to A.D. (R01 GM074215), to K.D.C. (R35 GM144121), to K.O. (R01 GM074207) and to P.L-G. (R35 GM150786). J.H. was supported by an NSF Graduate Research Fellowship during the early phases of this project (grant #1650112). A.D. and K.O. acknowledge partial salary support from the Ludwig Institute for Cancer Research.

## AUTHOR CONTRIBUTIONS

TK made initial observations that established the project and built the somatic degradation system for BUB-1. JH designed and conducted the majority of experiments with guidance from PLG. HH created the semi-automated CellProfiler image analysis pipeline. CV and EC conducted experiments with guidance from JH. KDC helped design TPR mutants based on structural homology. A. Deep and KDC generated and helped interpret AlphaFold models of the BUB-1 TPR domain interfacing with KNL-1. JH and A. Desai designed figures and wrote the manuscript, incorporating feedback from all authors. KO and A. Desai obtained funding and supervised the project.

## MATERIALS AND METHODS

### Worm Strains

*C. elegans* strains used in the study are listed in **Table S1** and were maintained at 20°C. RNAi-resistant *bub-1* and *knl-1* transgenes were previously described (Espeut et al., 2012; Moyle et al., 2014). RNAi-resistant *bub-3* transgenes were made by re-encoding the second exon of *bub-3* (**Figure S4B**). The first intron of *bub-3* was shortened to remove DNA hairpin secondary structure. The *bub-3* deletion strain has been previously described (Kim et al 2015).

For ZF1::GFP::BUB-1, the PIE-1 CCCH type zinc-finger domain (ZF1; 97-132 aa) was amplified from genomic DNA and inserted after the start codon. Then, the SPGGGGGG linker, GFP, and *bub-1* sequences were fused, inserted into pCFJ151, and injected into strain EG6429. For ZF1 (mutant)::GFP::BUB-1, mutations C103S and C113S were introduced in the PIE-1 CCCH type ZF1 (97-132aa) region.

GFP with introns enriched for Periodic An/Tn Clusters (GFP(PATC enriched)) was obtained from the Frokjaer-Jensen lab (Aljohani et al., 2020). All transgenes were cloned into plasmids pCFJ151 or pCFJ352 and injected into strains EG6429 or EG6701, respectively, along with plasmids encoding for the Mos transposase and a mix of four negative selection markers to remove extrachromosomal plasmid arrays. Successful integrants were selected by rescue of the *unc* phenotype of the parental strains and by a lack of negative selection markers 7-10 days after injection. Integrations were confirmed by PCR.

*In-situ* GFP tagged *bub-1* and *knl-1* were previously described (Cheerambathur et al 2019). The *in-situ* GFP tagged *bub-3* strain was created by plasmid based CRISPR Cas9 mutagenesis (Waaijers et al., 2013). In brief, plasmids containing Cas9 and sgRNA targeting the end of the *bub-3* coding sequence, along with a plasmid containing the *bub-3::gfp* repair template with 1000 base pair homology arms upstream and downstream of the *bub-3* stop codon and two co-injection markers, were injected into the germlines of N2 adult worms. The sgRNA sequence was AATTATTTCGGTCTGCTCTC. Fluorescent co-injection markers pGH8 and pCFJ90 form extrachromosomal arrays and express mCherry in the pharynx and neurons. F1 progeny expressing the fluorescent co-injection markers were singled out, and allowed to lay progeny for one week. After one week, the plates were washed and used to generate worm lysate for PCR-based genotyping to confirm integration of the GFP tag.

### RNAi

dsRNAs used in the study are listed in **Table S2** and were synthesized *in vitro*. Target sequences were amplified by PCR using oligos containing either T3 or T7 promoters at their 5’ end. MEGAscript T3 or T7 RNA polymerases (Thermofisher) were used to synthesize complementary RNAs from PCR templates. The MEGAclear kit (ThermoFisher) was used to purify the single-stranded RNA products. Single-stranded RNA was annealed by incubation at 68°C for 10 minutes followed by 37°C for 30 minutes. The concentration of the resulting dsRNA were measured by A260nm using a NanoDrop spectrophotometer (Thermo Fisher Scientific).

For RNAi-based gene knockdowns, 1 mg/mL of dsRNA was injected into L4 stage worms. Injected worms were incubated at 20°C for 36-48 hours before imaging-based experiments. For embryonic viability assays, L4 stage worms were injected with dsRNA and recovered at 20°C for 24 hours. The injected worms were singled and laid progeny for 24 hours at 20°C. Parental worms were then removed and after a further 24 hours, progeny was scored as either viable or dead.

### Fluorescence imaging of *C. elegans* embryos

Gravid adult hermaphrodite worms were dissected in M9 buffer, early embryos were transferred onto a 2% agarose pad using a mouth pipet, covered with a 22x22 mm coverslip, and imaged at 20°C. To assay kinetochore recruitment, embryos were imaged on an Andor Revolution confocal system (Andor) coupled to a CSU-10 spinning disk confocal scanner (Yokogawa) and an electron multiplication back-thinned charge-coupled device camera (iXon, Andor), using a 100x 1.4 NA Plan Apochromat objective; or with an inverted Axio Observer Z1 (Zeiss) microscope with a CSU-X1 spinning disk head (Yokogawa) and a 100x 1.3 NA Plan Apochromat Lens (Zeiss). A 6x2 μm z-stack was collected every 10 or 20 seconds.

For assaying mitotic timing and chromosome dynamics, one-cell embryos expressing GFP::H2b were imaged on a widefield deconvolution microscope (DeltaVision) connected to a charge-coupled device camera (pco.edge 5.5 sCMOS; PCO) and a 60x 1.42NA PlanApo N objective (Olympus). A 5x2 μm z-stack was collected every 10 seconds.

### Imaging analysis and quantifications

Microscope images were processed using ImageJ. NEBD was defined as the time point where free nuclear histone signal dissipates into the cytosol and spindle forces start pushing on chromosomes. Anaphase was the first time point where the separation of sister chromatids was evident. For kinetochore recruitment assays, a box was drawn around the kinetochores, and the integrated density was recorded. A second box was drawn 5 pixels wider than the first in order to subtract out the local background.

For measuring BUB-1 dynamics, CellProfiler version 3.1.9 software (McQuin et al., 2018) was used to quantify kinetochore localization of GFP tagged proteins over time (**Fig S4A)**. In brief, maximum intensity projections were made for each hyperstack in ImageJ. Projected movie files were imported into CellProfiler for analysis. A Laplacian of Gaussian function was used to segment chromosomes in the mCh::H2b channel. Co-localized GFP signal was then quantified in the segmented area. The segmented object was expanded by three pixels to subtract out local background. The process was repeated for each frame in an image sequence to generate dynamic curves over time. Data were normalized to the peak value of the wildtype.

### Protein expression and purification

6xHis tagged *C. elegans* KNL-1 (1-505) wildtype and mutant constructs were cloned into pET21a vectors and transformed into Rosetta BL21(DE3) pLysS *E. coli* cells. Cultures were grown at an OD600 of 0.7-0.8 and induced overnight at 18°C with 0.3 mM IPTG. The next morning, cells were collected by centrifugation, washed with chilled 1x PBS, and resuspended with resuspension buffer (25 mM Tris, pH 7.5, 5 mM Imidazole, 300 mM NaCl, 5 mM B-ME, 10% glycerol) supplemented with 1 mM PMSF, 2 mM benzamidine, cOmplete Protease Inhibitor, and 4 units/mL DNAseI. Cells were lysed by sonication, and lysates were cleared by ultracentrifugation in a Ti45 rotor at 28,000 rpm for 30 minutes at 4°C. 6xHis tagged proteins were purified from the clarified lysates by using Ni-NTA beads (Qiagen), and incubated with end-over-end rotation overnight at 4°C. Beads were washed twice with re-suspension buffer and once with 20 mM imidazole in resuspension buffer before applying to polypropylene columns (Qiagen). Samples were eluted in 1 mL fractions using 400 mM Imidazole in resuspension buffer. Fractions containing the most protein were combined and dialyzed against storage buffer (20 mM Tris-Cl, pH 7.5, 150 mM NaCl, 1 mM MgCl2, 5 mM βME, 0.1% Triton X-100, 5% glycerol). Proteins were concentrated using 10,000 NMWL filter (Amicon), and the concentration was assessed by BSA standard curve on a Coomassie gel. Proteins were aliquoted, snap frozen in liquid nitrogen, and stored at -80°C.

### In vitro *binding* assays

A construct encoding for *C. elegans* BUB-1 TPR (1-189)::3xFLAG under a CMV promoter was transfected into Freestyle 293-F cells (Thermo Fisher Scientific). 48 hours later, cells were harvested and washed with PBS before re-suspending in lysis buffer (20 mM Tris-Cl pH 7.5, 50 mM NaCl, 5 mM EGTA, 2 mM MgCl2, 0.5% Triton X-100, 1 mM DTT), supplemented with 1x cOmplete Protease inhibitor cocktail (Millipore). Cells were sonicated for 6 minutes in an ice-cold water bath, then centrifuged for 15 minutes at 15,000xg at 4°C. Cell lysates were incubated with M2 anti-FLAG beads (Sigma) while rotating for 2 hours at 4°C. After the two hour incubation, beads were washed four times with lysis buffer. Meanwhile, 2 μM KNL-1::6xHis (1-505) was phosphorylated with 1 μM constitutively active PLK-1 T194D (gift from Jeffrey Woodruff, UT Southwestern) in kinase buffer (20 mM Tris-Cl pH 7.5, 50 mM NaCl, 10 mM MgCl2, 1 mM DTT, 0.2 mM ATP), for 2 hours at 23°C. Phosphorylated KNL-1::6xHis was then directly added to BUB-1 TPR::3xFLAG bound to beads to a final KNL-1::6xHis concentration of 100 nM. Binding was performed with rotation for 2 hours at 4°C, after which beads were washed with lysis buffer containing 0.05% Triton X-100 4 times. Proteins were eluted from the beads by resuspending in SDS sample buffer before analysis by immunoblotting.

### Preparation of worm lysates

L4-stage worms were incubated at 20°C for 36-48 hours. 60 worms per condition were washed with M9 plus 0.1% Tween 20, resuspended in SDS-sample buffer and lysed by sonication followed by boiling.

#### Immunoblotting

Antibodies used in this study are listed in **Table S3**. For binding assays, samples were run on 4-15% gradient Mini-Protean gels (BioRad) and transferred to PVDF membranes using a Transblot Turbo system (BioRad). The membranes were incubated in blocking buffer (5% skim milk in TBS-Tween 0.05%) for one hour, and antibodies were incubated in either the same buffer or in TBS-T + 5% BSA. Membranes were developed using Western Bright Sirius substrate (Advansta) before imaging on a ChemiDoc system (BioRad).

For blots of worm lysates, 4-12% NuPAGE Bis-Tris gels (Invitrogen) were loaded with lysate equal to 5-10 adult worms. Proteins were transferred overnight to nitrocellulose membranes, after which membranes were blocked and antibodies incubated in TBS-T with 5% milk. Membranes were developed using Western Bright Sirius substrate (Advansta) before imaging on ChemiDoc System (BioRad).

### AlphaFold2 Structure Prediction

Structure prediction for BUB-1 (residues range: 1-190) and KNL-1 (residues range: 1-500) was performed using AlphaFold2 ColabFold notebook (Jumper et al., 2021; Mirdita et al., 2022). To model the ternary complex of KNL-1, BUB-1, and BUB-3 proteins, the sequences of full-length BUB-3, the N-terminal region of BUB-1 containing the TPR and BUB-3 binding domain (aa 1-255 of BUB-1), a MELT1-containing peptide (aa 74-96 of KNL-1), and a TF1-containing peptide (aa 116-132) were included as separate chains. The top-scoring model was selected based on ranking and comparison with the *S. cerevisiae* Bub3-Bub1-pMELT crystal structure (PDB ID 4BL0). Structural analysis and depictions were carried out in PyMol (DeLano, 2002) and Chimera X (Meng et al., 2023).

### Phylogenetic Analysis

Sequences of *knl-1* genes across nematode species were downloaded from Wormbase website. Initial motif discovery was performed using the MEME algorithm (Bailey et al., 2009); this was followed up by manual annotation of sequences with D/E/N residues in the -2 position, T in the 0 position, and F in the +4 position. Cladogram was created using EMBL Simple Phylogeny tool (Madeira et al., 2022). Secondary structure of KNL-1 in different nematode species was predicted using PsiPred (Jones, 1999).

#### Statistical analyses

Unpaired t tests were used to calculate p values in Prism software (Graphpad). P values are displayed as follows: ns = p>0.05; ** = p<0.01; *** = p<0.001; **** = p < 0.0001. Error bars display the 95% confidence interval.

---

**Supplemental Figure 1:**
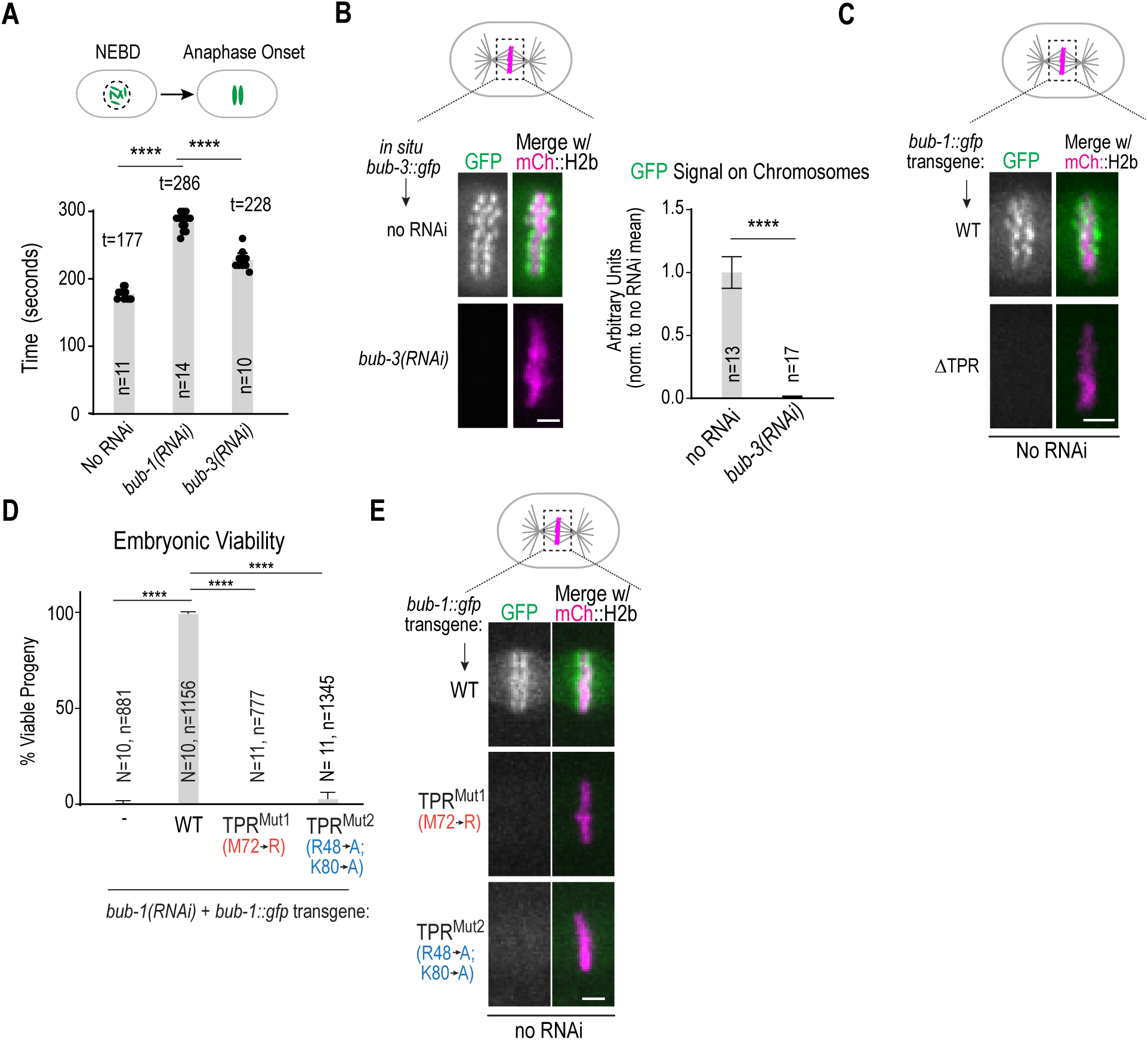
Penetrance of BUB-3 depletion and additional analysis of BUB-1 TPR mutants. **(A)** The interval from nuclear envelope breakdown (NEBD) to anaphase onset was measured in the indicated conditions; *n* refers to the number of embryos imaged. **(B)** (*left)* One cell embryos expressing *in situ* GFP-tagged BUB-3 were imaged in the indicated conditions. Scale bar is 2 µm. (*right)* Quantification of BUB-3::GFP kinetochore localization indicating that *bub-3(RNAi)* is highly penetrant. Error bars are the 95% confidence interval. **(C)** Images of aligned chromosomes in one cell *C. elegans* embryos expressing the indicated *bub-1::gfp* transgenes in the presence of endogenous BUB-1 (No RNAi). ΔTPR BUB-1 fails to localize even in the presence of endogenous BUB-1. Scale bar is 2 µm. (**D)** Embryonic viability analysis for the indicated conditions. *N* is the number of worms and *n* the number of progeny embryos scored. Error bars are the 95% confidence interval. **(E)** Images of aligned chromosomes in one cell *C. elegans* embryos expressing the indicated *bub-1::gfp* transgenes in the presence of endogenous BUB-1 (No RNAi). Both TPR^mut1^ and TPR^mut2^ BUB-1 fail to localize even when endogenous BUB-1 is present. Scale bar is 2 µm. All *p* values were calculated from unpaired t tests; **** = p<0.0001.

**Supplemental Figure 2:**
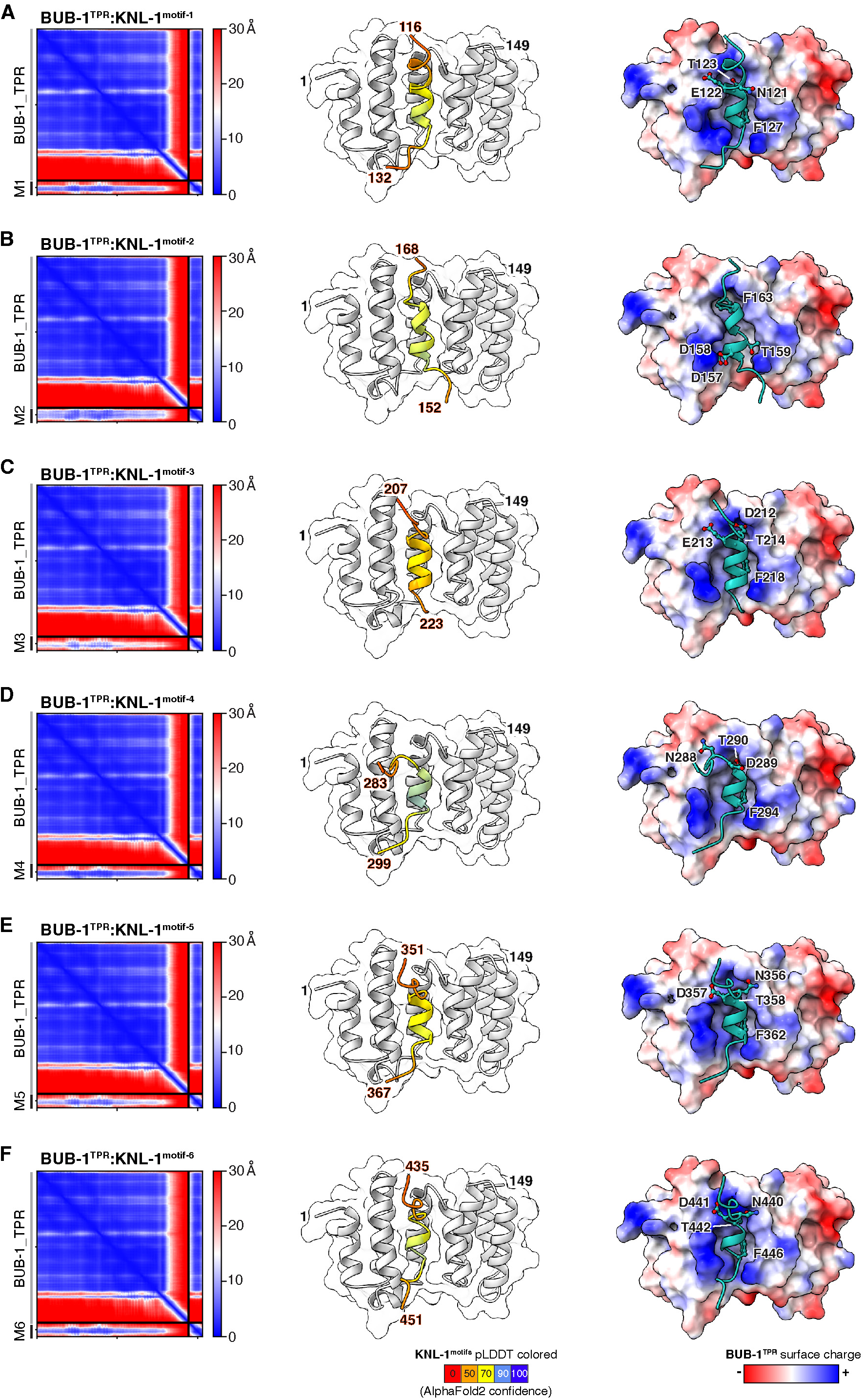
AlphaFold models of the BUB-1 TPR with the 6 TF motifs in the KNL-1 N-terminus. **(A) – (F)** AlphaFold models of BUB-1 TPR (aa 1-149) interfacing with TF motifs 1-6 (*panels A-F*) in the KNL-1 N-terminus. For each model, 3 elements are shown: *(left*) Predicted Aligned Error (PAE) plot; (*middle*) cartoon model showing the interface of the TPR and the TF motif; the TF motif is color-coded based on the predicted local distance difference test (pLDDT) score; (*right*) BUB-1 TPR surface charge depiction with the modeled bound TF motif; specific residues of the TF motif are highlighted. Aside from motif 2, the F residue of the TF motif occupies the same hydrophobic pocket in the TPR domain. The basic character in the vicinity of the T residue, targeted by PLK-1, likely accounts for the phospho-dependence of the interaction.

**Supplemental Figure 3:**
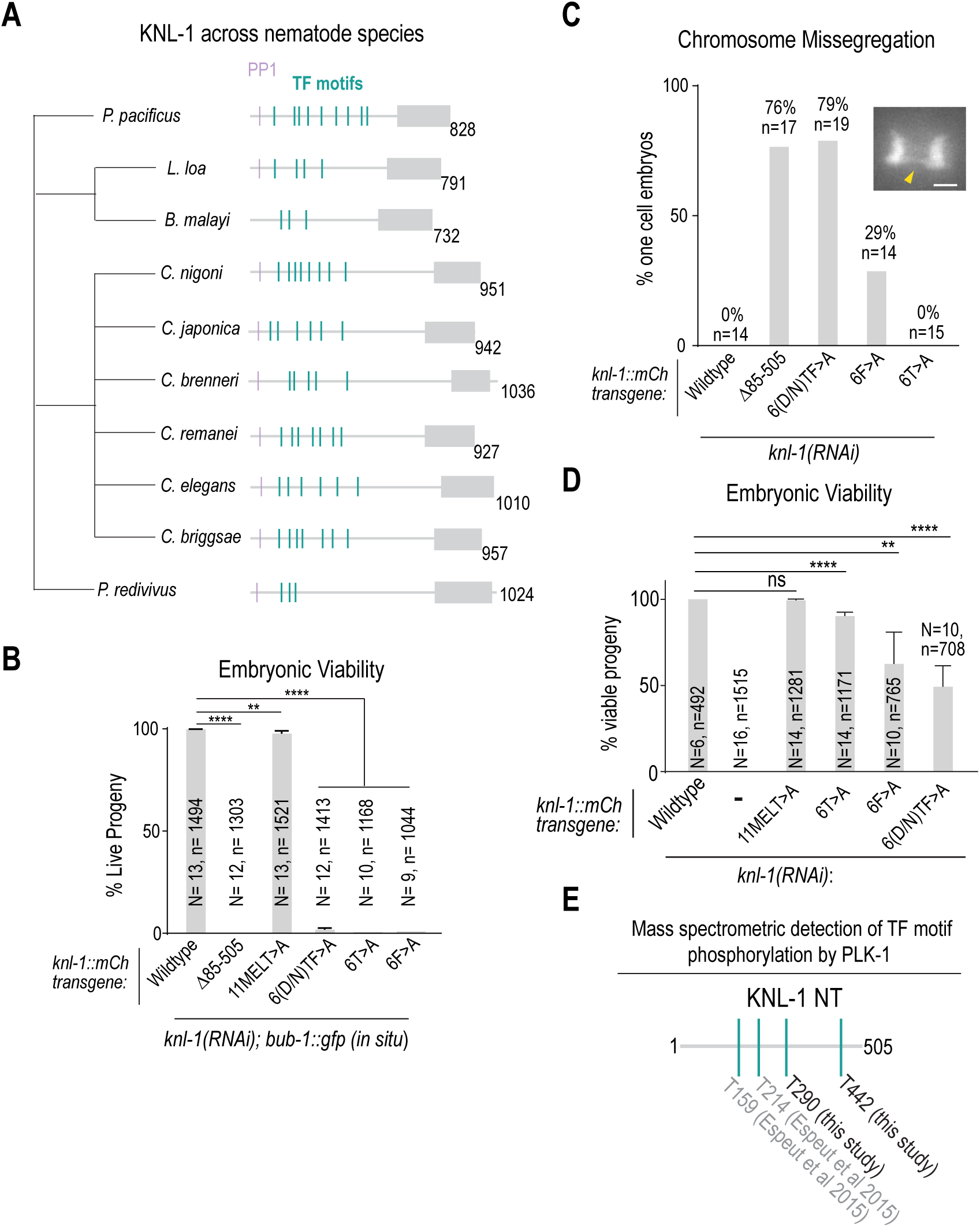
Additional analysis of KNL-1 TF motifs. **(A)** Cladogram showing TF motifs present in *knl-1* genes across different nematode species. **(B)** Embryonic viability was scored for indicated conditions; note that this analysis was conducted in the presence of *in situ* GFP-tagged BUB-1. *N* is the number of worms and *n* is the number of progeny scored. Error bars are the 95% confidence interval. **(C) & (D)** Chromosome missegregation and embryonic lethality analysis for the indicated KNL-1 variants when the endogenous *bub-1* locus was untagged. *(C)* Chromosome missegregation was quantified for each of the indicated conditions. *n* is the number of embryos imaged. Inset image shows example of missegregating chromosome, highlighted by yellow arrowhead. Scale bar of inset is 2 μm. Example image is the same as shown in Fig. 1B and Fig. 2H. *(D)* Embryonic viability quantified for indicated conditions. *N* is the number of worms and *n* is the number of progeny scored. Error bars are the 95% confidence interval. **(E)** Schematic of TF motif phosphorylation by PLK-1 detected by mass-spectrometry. Phosphorylation of T290 and T442 (TF motifs 4 and 6 respectively) was identified in this study. Phosphorylation of T159 and T214 (TF motif 2 and TF motif 3 respectively) was identified in a previous study (Espeut et al., 2015). All p values were calculated from unpaired t tests; ns = not significant, ** = p< 0.01, **** = p<0.0001.

**Supplemental Figure 4:**
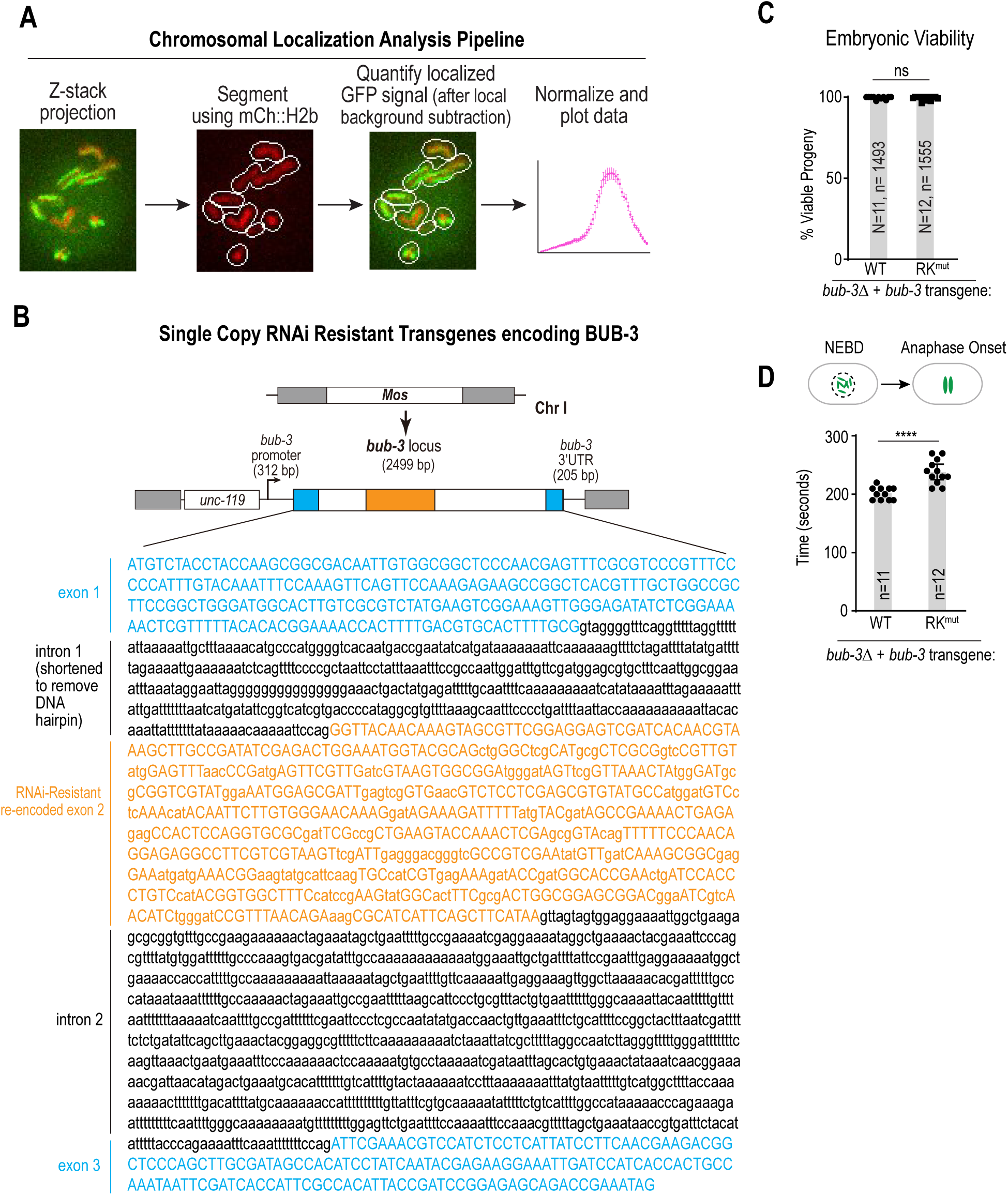
Image analysis approach to quantify chromosomal localization, design of the RNAi-resistant bub-3 transgene and additional phenotypic analysis of RK^mut^ BUB-3. **(A)** Overview of CellProfiler-based analysis to segment chromosomes based on the mCh::H2b signal and measure chromosomal GFP fluorescence. A Laplacian of Gaussian function was applied to the maximum intensity projection of mCh::H2b images in the timelapse sequence to identify chromosomes and expand a region around them. The GFP signal within the segmented regions was quantified, and the region was further expanded by three pixels to subtract local background. The process was applied to each timepoint to generate dynamic chromosomal localization curves for each condition. **(B)** Schematic of RNAi-resistant *bub-3* transgenes. Exon 2 was reencoded to preserve coding information but make the transgene-encoded mRNA resistant to RNAi triggered by a dsRNA raised to the endogenous sequence. In addition, intron 1 was shortened to remove a DNA hairpin. **(C)** Embryonic viability was quantified for each of the indicated conditions. **(D)** The interval from NEBD-Anaphase onset was quantified for each of the indicated conditions in one cell *C. elegans* embryos. *n* refers to the number of embryos imaged. All p values were calculated by unpaired t tests; ns = not significant, **** = p<0.0001.

**Table S1:**
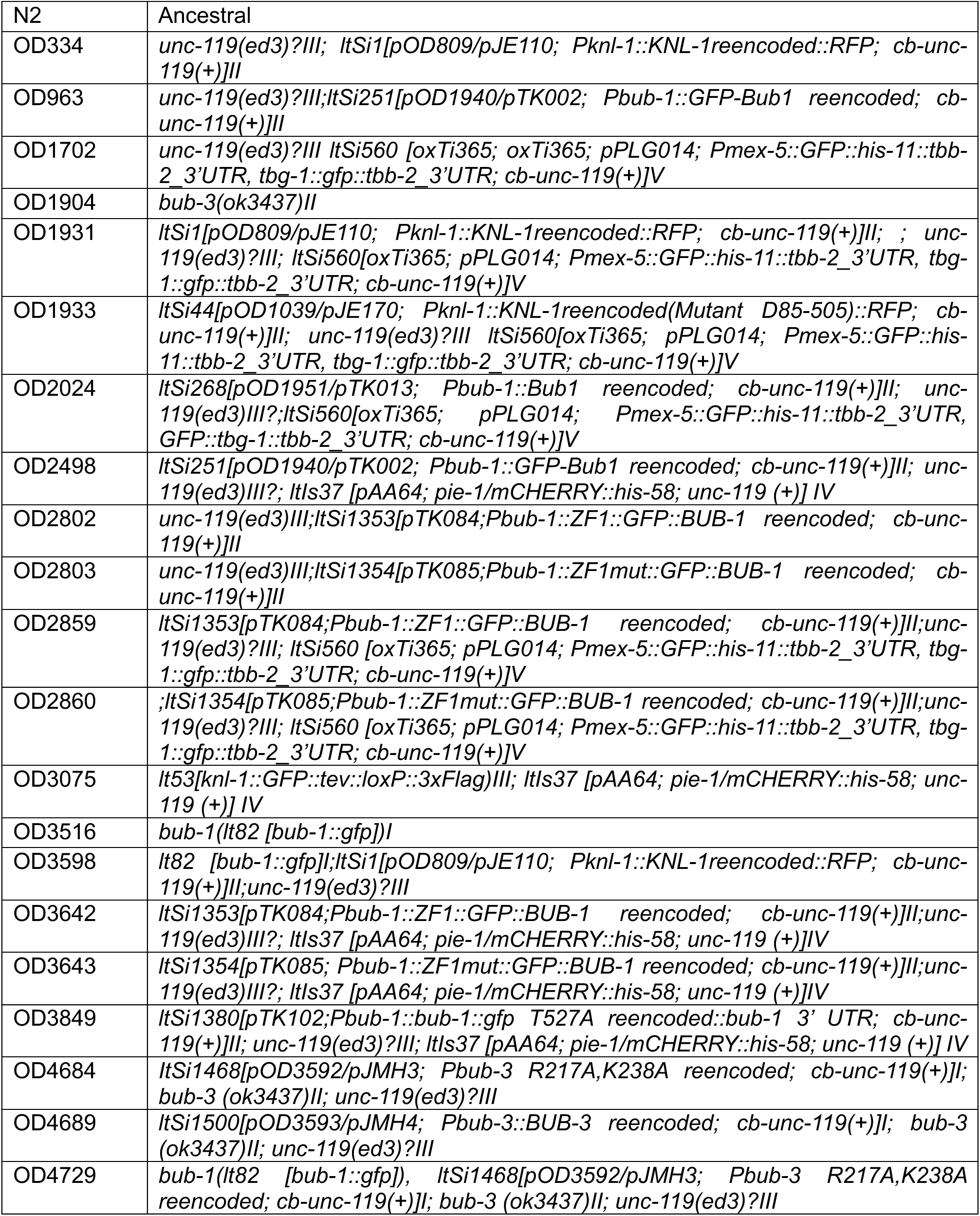
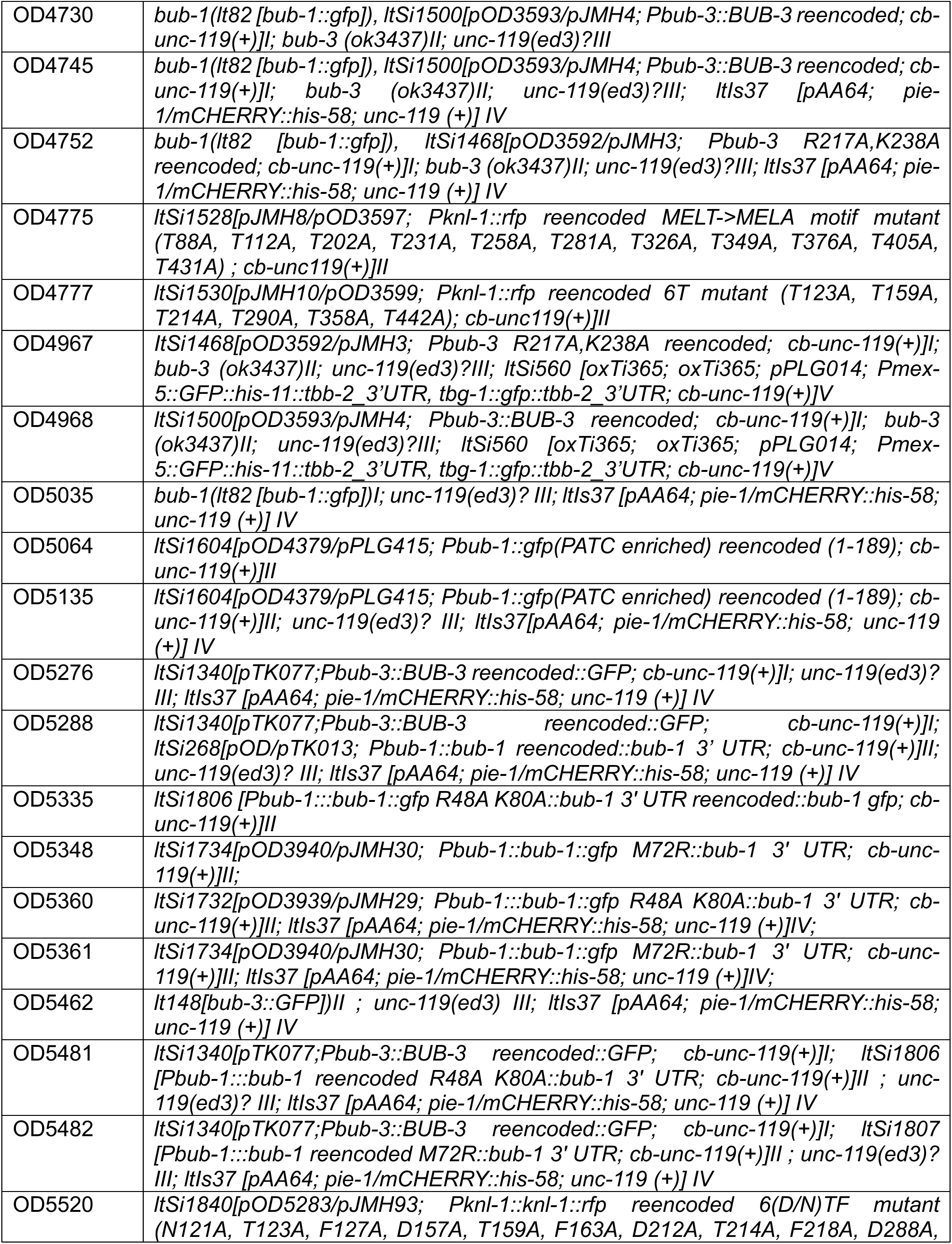
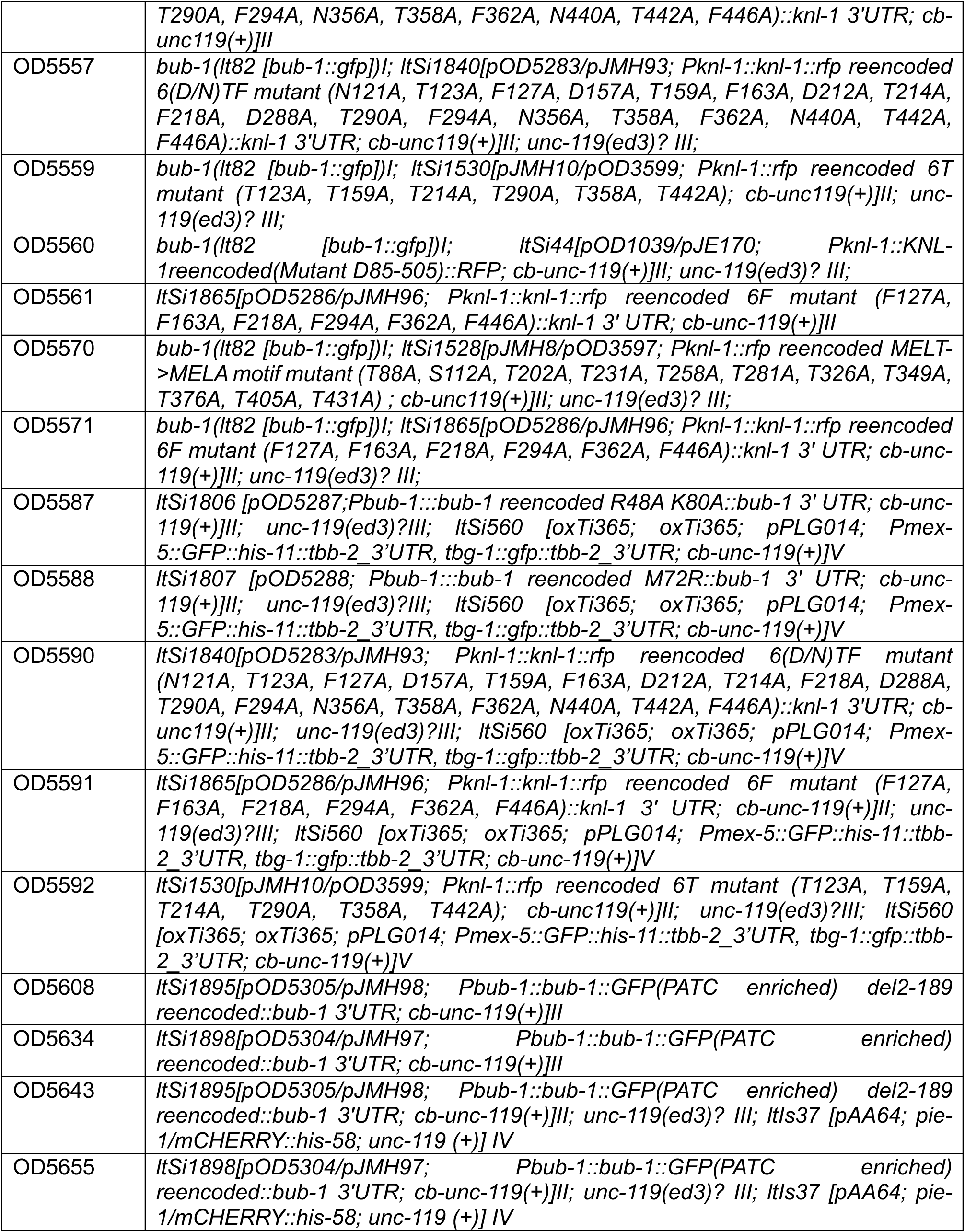
*C. elegans* Strains.

**Table S2:**
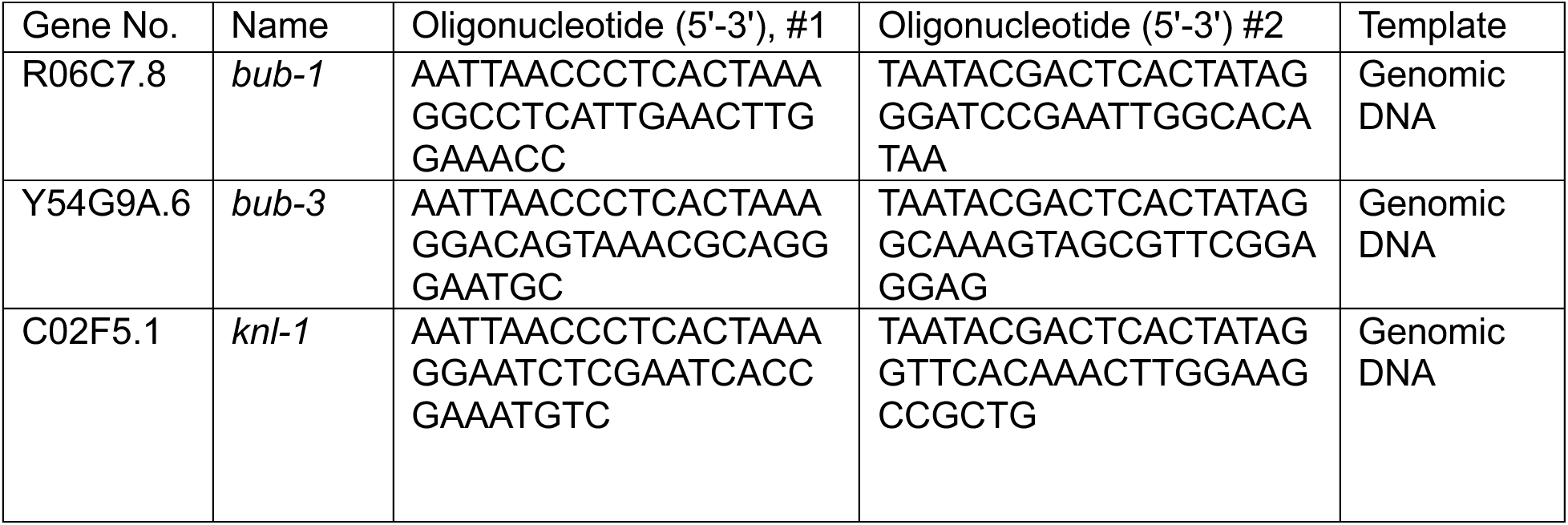
Primers for dsRNA synthesis.

**Table S3:** Antibody list.

| Antibody Name | Target | Host Species | Stock Concentration (mg/mL) | Source |
| --- | --- | --- | --- | --- |
| OD31-B | BUB-1 (aa 287-665) | Rabbit | 2.8 | Oegema et al. 2001 |
| OD194 | GFP | Goat | 15 | Hyman Lab |
| DM1α | α-tubulin | Mouse | 1 | Sigma (T9026) |
| OD144 | BUB-3 | Rabbit | 2.98 | Essex et al, 2009 |
| M2 | FLAG | Mouse | 1 | Sigma (F1804) |
| OD34 | KNL-1 (1-150) | Rabbit | 3.8 | Desai et al 2003 |
|  | Anti-Rabbit IgG, HRP conjugated | Goat | 0.8 | Jackson Immunoresearch (111-035-003) |
|  | Anti-Mouse IgG, HRP conjugated | Donkey | 0.8 | Jackson Immunoresearch (715-035-150) |
|  | Anti-Mouse Heavy Chain Specific, HRP conjugated | Goat | 0.8 | CST (96714) |
|  | Anti-Rabbit IgG, HRP conjugated | Donkey | 0.8 | Jackson Immunoresearch (711-035-152) |
|  | Anti-Goat IgG, HRP conjugated | Donkey | 0.5 | Jackson Immunoresearch (705-035-003) |
|  | Anti-Mouse IgG, AP conjugated | Donkey | 0.8 | Jackson Immunoresearch (715-055-150) |

